# Ventral Striatum is Preferentially Correlated with the Salience Network Including Regions in Dorsolateral Prefrontal Cortex

**DOI:** 10.1101/2024.10.13.618063

**Authors:** Heather L. Kosakowski, Mark C. Eldaief, Randy L. Buckner

## Abstract

The ventral striatum (VS) receives input from the cerebral cortex and is modulated by midbrain dopaminergic projections in support of processing reward and motivation. Here we explored the organization of cortical regions linked to the human VS using within-individual functional connectivity MRI in intensively scanned participants. In two initial participants (scanned 31 sessions each), seed regions in the VS were preferentially correlated with distributed cortical regions that are part of the Salience (SAL) network. The VS seed region recapitulated SAL network topography in each individual including anterior and posterior midline regions, anterior insula, and dorsolateral prefrontal cortex (DLPFC) – a topography that was distinct from a nearby striatal seed region. The region of DLPFC linked to the VS is positioned adjacent to regions associated with domain-flexible cognitive control. The full pattern was replicated in independent data from the same two individuals and generalized to 15 novel participants (scanned 8 or more sessions each). These results suggest that the VS forms a cortico-basal ganglia loop as part of the SAL network. The DLPFC is a neuromodulatory target to treat major depressive disorder. The present results raise the possibility that the DLPFC may be an effective neuromodulatory target because of its preferential coupling to the VS and suggests a path toward further personalization.

---

The basal ganglia interact with the cerebral cortex via multiple partially segregated circuits (Alexander, DeLong, and Strick 1986; Haber, 2016; Heimer, Switzer, and Van Hoesen 1982). The ventral striatum (VS), which includes the nucleus accumbens (NAc) and has been associated with reward and motivation, receives inputs from distributed cortical regions and projects back to the cortex through the thalamus (Alexander, DeLong, and Strick 1986; Haber and McFarland 1999; Heimer 1978). Multiple regions in orbitofrontal, cingulate, temporal, and insula cortices project to the VS (Chikama et al. 1997; Haber et al. 1995; Haber, Fudge, and McFarland 2000; Kunishio and Haber 1994; Selemon and Goldman-Rakic 1985). Additionally, subcortical structures including the hippocampus, ventral tegmental area (VTA), and subnuclei of the amygdala project to the VS suggesting a complex brain-wide organization that is broadly associated with ‘limbic’ functions in addition to its role in reward circuitry (Brog et al. 1993; Fox 1943; Groenewegen et al. 1987; Haber et al. 2000; Lynd-Balta and Haber 1994; for reviews, see Groenewegen et al. 1999; Heimer 2003). The details of projection patterns to the VS, including understanding if the VS is interacting preferentially with specific cortical networks, have translational implications for the treatment of addiction and mood disorders (e.g., Drevets, Price, and Furey, 2008; Koob and Volkow, 2016).

In a particularly thorough investigation, Choi, Haber and colleagues (2017) compiled a series of cases of both cortical anterograde tracer injections and retrograde injections directly to the subdivisions of the VS. Critically, they examined the injection patterns in relation to separate ventral (VSv) and dorsal (VSd) zones demarcated by acetylcholinesterase (AChE) staining. VSv includes the NAc shell, and VSd includes the NAc core extending into the ventral portions of the caudate and putamen. They found that the cortical projections to the striatum differed as the region moved from the most ventral portions of the VS (VSv) medially into the caudate (medial VSd) and laterally into the putamen (lateral VSd). The projection patterns did not respect the estimated AChE border but robust differences in connectivity were nonetheless observed. Of particular interest, the VSv received extensive projections from orbitofrontal and medial temporal cortex that were convergent with dense projections from the basolateral amygdala. These findings raise the possibility that differences in extrinsic connectivity may form segregated (or partially segregated) networks within and around the VS extending into the ventral caudate and putamen.

The organization of the striatum has been extensively studied in humans using functional connectivity. Functionally connectivity MRI (fcMRI) estimates connectivity based on the correlated fluctuations in the blood oxygenation level-dependent (BOLD) signal and is an indirect, imperfect proxy for anatomical connectivity (for discussions and caveats see Buckner, Krienen, and Yeo 2013; Fox and Raichle 2007; Murphy, Birn, and Bandettini 2013; Power, Schlaggar, and Petersen 2014; Smith et al. 2013; Van Dijk et al. 2010). Nonetheless, estimates of striatal connectivity with the cortex have been reliable and broadly converge with monkey anatomical observations, including distinctions between motor, association, and limbic zones (Choi, Yeo, and Buckner 2012; Di Martino et al. 2008; Greene et al. 2014; Jarbo and Verstynen 2015; Marquand, Haak, and Beckmann 2017; Morris et al. 2016). However, due to challenges specific to studying the human VS, a detailed account of its organization has been difficult to unravel.

First, the VS is anatomically heterogenous. Rodent studies indicate there are fine distinctions and cell classes that possess specificity at a resolution beyond what human techniques can presently observe. However, the extrinsic connectivity differences that have been noted in monkeys suggest that adjacent subregions may be connected to distinct cortical networks. Notably, moving from the ventral portions of the VS dorsally into the caudate and putamen (what Choi, Ding, and Haber 2017 refer to as VSd) produces different patterns of extrinsic connectivity. These coarse macroscale anatomical differences are at a spatial resolution that fcMRI should be able to capture.

An additional challenge is that the VS is near to regions that have susceptibility artifacts (Ojemann et al. 1997). This necessitates careful attention to signal properties. In slice acquisitions commonly used for fcMRI, the VS is adequately sampled, as it is positioned above the major portions of the susceptibility artifacts arising from the sagittal sinus, but the signal can nonetheless be less robust than other regions (see Figure 1 in Kosakowski et al. 2024). Collection of repeat acquisitions to boost signal-to-noise (SNR) properties, acquisition of single-echo or multi-echo acquisitions with short echo times (TE) (e.g., Kundu et al. 2017), careful attention to the signal quality and distortion around the VS, and replication and generalization are all paths to mitigate challenges related to susceptibility artifacts.

**Figure 1.**
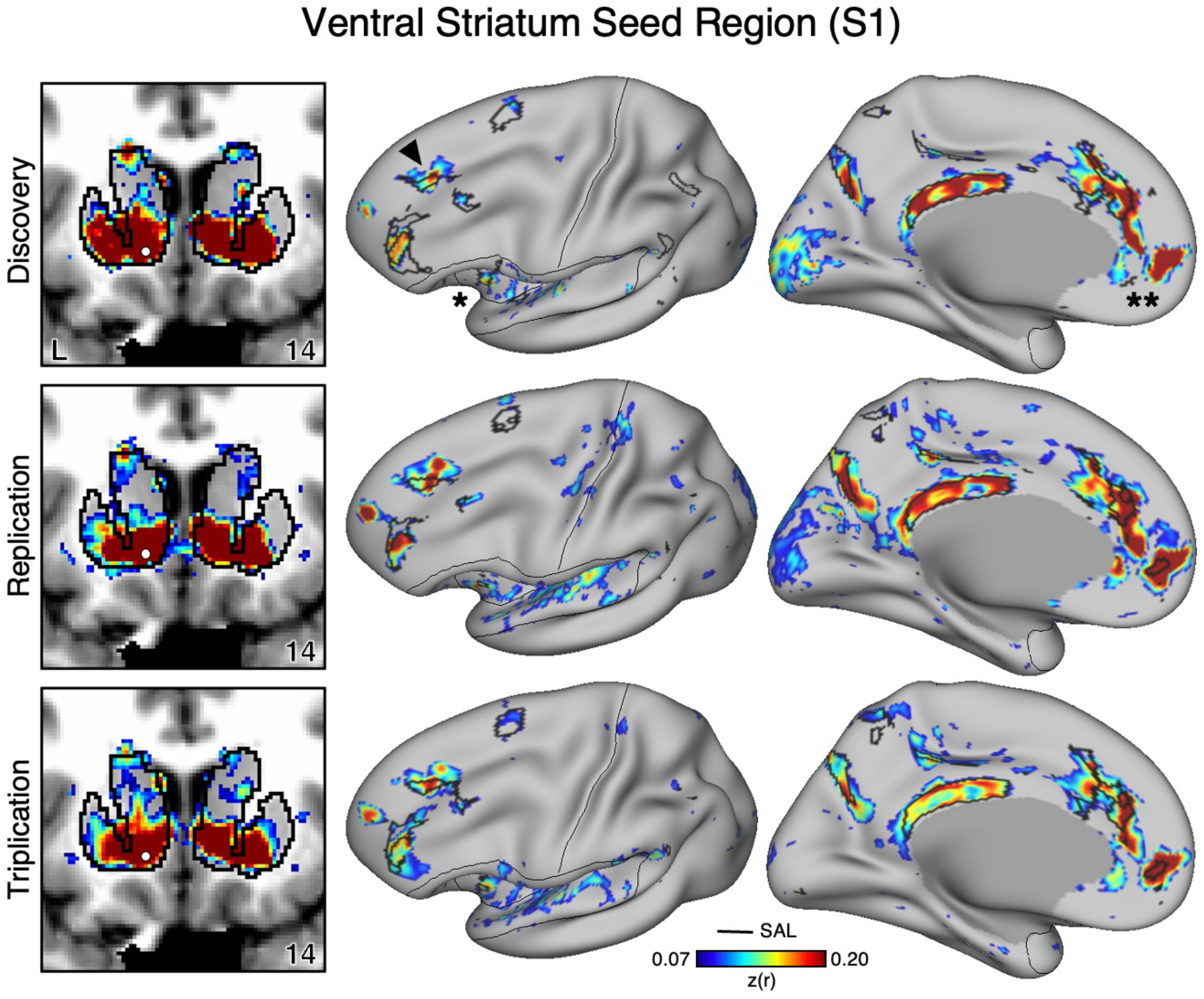
The Ventral Striatum (VS) Correlates with the Full Cortical Extent of the Salience (SAL) Network in S1. A seed region (indicated with a white circle) in the VS (top left) recapitulates the full extent of the SAL network, including the dorsolateral prefrontal cortex (indicated with a black arrow), insula (indicated by asterisk), anterior cingulate cortex, and rostral portions of the medial prefrontal cortex (indicated by double asterisks). In independent data from S1, this pattern was replicated (second row) and triplicated (third row) using the same seed region (white circles). Cortical results are visualized on the inflated fsaverage6 surface with the lateral view slightly tilted to show the dorsal surface. Striatal results are visualized in coronal sections on each individual’s own T1-weighted anatomical image transformed to the atlas space of the Montreal Neurological Institute (MNI) atlas. L = left. Coordinates in the bottom right of the left panels indicate the y coordinates (in mm). Black outlines in the volume indicate the boundaries of the striatum and outlines on the cortical surface indicate the SAL network (see methods). All correlation values are plotted after r to z conversion using the jet colorscale shown by the legend.

A final challenge is that the proximity of VS and cortical regions can lead to signal blur at low spatial resolutions and especially when employing group averaging. Blur from the cortex was mitigated in the original comprehensive maps of the striatum in Choi et al. (2012) using signal regression. However, in retrospect, the maps associated with the VS in this earlier report likely contained residual artifacts of spatial blur. Specifically, correlation patterns for seed regions placed in the VS recapitulated the cortical network commonly known as the ‘Default’ network (see Figure 11 in Choi et al. 2012; see also the VSi and VSs seed correlation maps in Di Martino et al. 2008) including correlation with the subgenual anterior cingulate (sgACC). sgACC is a cortical region located on the frontal midline adjacent to the VS. Examining the correlation pattern at or near sgACC in group-averaged data, for example using the openly available neurosynth platform, yields a pattern that extends through orbitofrontal and ventral midline regions that are affected by susceptibility artifacts (see coordinates X = -4, Y = 16, Z = -8 in https://neurosynth.org/locations/; Yarkoni et al. 2011). These challenges can be partially surmounted by examining data within the individual.

By respecting idiosyncratic neuroanatomy within individuals, precision fcMRI has recently revealed additional details of VS organization (Gordon et al. 2022; Greene et al. 2020; Lynch et al. 2024). Greene and colleagues (2020) showed that distinct regions within and near to the VS correlate with distributed networks including the Salience (SAL) network as well with the ‘Default’ network in multiple individuals. The finding of correlations with the SAL network is particularly intriguing given the network’s proposed role in orienting to transient, relevant events (Seeley et al. 2007; Seeley 2019; Uddin 2015). Lynch and colleagues (2024) recently undertook a detailed assessment of striatal regions functionally correlated with the SAL network. In both a control cohort and separately in a group of individuals with Major Depressive Disorder (MDD), the SAL network was consistently associated with the VS extending into the ventral caudate. The effect was found in most individuals and generalized across both groups (see Figure 1e in Lynch et al. 2024).

Motivated by these recent discoveries, we employed precision approaches to revisit VS organization. To do so, we first analyzed fMRI data from two intensively scanned individuals (data collected over 31 sessions each). These data enabled us to estimate VS functional connectivity, respecting idiosyncratic anatomical variability, and replicate the results within the same participants. We next generalized the findings to 15 additional participants (data collected over 8 or more sessions each). In all analyses the SAL network was coupled to the VS, replicating Lynch et al. (2024). Adjacent regions in the ventral putamen were coupled to a distinct network often referred to as the Cingulo-Opercular (CG-OP) network (see also Gordon et al. 2023). Of particular interest, the pattern of correlations to the VS included a specific region of dorsolateral prefrontal cortex (DLPFC) that is spatially variable across individuals and distinct from regions involved in domain-flexible cognitive control. The finding that a region in DLPFC is reliably associated with the VS but variably localized has practical implications for targeted neuromodulation.

## Methods

### Participants

Two participants labeled S1 and S2 were each scanned across 31 separate MRI sessions (ages 22 and 23 yrs, both women). Fifteen additional participants labeled P1-P15 were each scanned across 8 or more MRI sessions (mean age = 22.1 yrs, SD = 3.9 yrs, 9 women). All participants were right-handed, native English speakers with no history of severe neurological and psychiatric illness. Participants provided informed consent using protocols approved by the Institutional Review Board of Harvard University. MRI data for P1-P15 are openly available through the NIH repository (https://nda.nih.gov) and have been previously reported in papers focused on the cerebral cortex (Du et al. 2024), caudate (Kosakowski et al. 2024), and cerebellum (Saadon-Grosman et al. 2023). The data were reanalyzed here with a focus on the VS. Initial discoveries were made in S1 and S2, and then P1-P15 were used to generalize the findings.

### MRI Data Acquisition

Acquisition details have been previously described (Braga et al. 2019; Du et al. 2024; Kosakowski et al. 2024; Saadon-Grosman et al. In Press) and key details are repeated here. Data were acquired at the Harvard Center for Brain Science using a 3T Siemens MAGNETOM Prisma^fit^ MRI scanner. Data were collected using the vendor-supplied 64-channel and 32-channel phased-array head-neck coils (Siemens Healthineers, Erlangen, Germany). Data from the two coils were combined as they have been previously (Du et al. 2024; Kosakowski et al. 2024; Saadon-Grosman et al. 2022). Foam and inflated padding were used to reduce head motion. Participants were instructed to remain still, awake, and look at a rear-projected display through a custom-built mirror attached to the head coil. During functional scanning, participants fixated a centrally presented plus sign (black on a gray background). The scanner room was illuminated, and eyes were video recorded using an Eyelink 1000 Plus with Long-Range Mount (SR Research, Ontario, Canada). Alertness was scored during each functional run.

S1 and S2 each participated in 31 MRI sessions with 61 to 62 resting-state fixation runs acquired in total per individual. Each session involved multiple resting-state fixation runs acquired using BOLD contrast functional MRI (fMRI; Kwong et al. 1992; Ogawa et al. 1992). A custom multiband gradient-echo echo-planar pulse sequence developed by the Center for Magnetic Resonance Research (CMRR) at the University of Minnesota (Van Essen et al. 2013; Xu et al. 2012, 2013; see also Setsompop et al. 2012) was used: voxel size = 2.4 mm, repetition time (TR) = 1,000 ms, TE = 32.6 ms, flip-angle = 64°, matrix 88 × 88 × 65, anterior-to-posterior (AP) phase encoding, multislice 5x acceleration. 65 slices were automatically positioned (van der Kouwe et al. 2008). Signal dropout was minimized by selecting a slice 25° from the anterior-posterior commissural plane toward the coronal plane (Mennes et al. 2014; Weiskopf et al. 2006). Each run lasted 7 min 2 sec (422 frames with 12 frames removed for T1 equilibration). No generalized auto-calibrating partial parallel acquisition (GRAPPA) acceleration was used which, when combined with multiband acceleration, can cause poor data quality (Wall 2023). A dual-gradient-echo B0 fieldmap was acquired to correct for spatial distortions: TE = 4.45 and 6.91 ms; slice prescription / resolution matched to the BOLD sequence.

P1-P15 each participated in 8-11 sessions with 15 to 24 resting-state fixation runs acquired in total per individual. BOLD acquisition parameters were similar to those used for S1 and S2 except the matrix was 92 × 92 × 65 (FOV = 221 × 221). For P12, the first two sessions were acquired with a different FOV (211 × 211); as such, the matrix was: 88 × 88 × 65, matching S1 and S2. Each fixation run was 7 min 2 sec (422 frames, first 12 removed for T1 equilibration).

For S1 and S2, at least one rapid T1w structural scan was collected for each participant using a multi-echo magnetization prepared rapid acquisition gradient echo (ME-MPRAGE) 3D sequence (van der Kouwe et al. 2008): voxel size = 1.2 mm, TR = 2,200 ms, TE = 1.57, 3.39, 5.21, 7.03 ms, TI = 1,100 ms, flip-angle = 7°, matrix 192 × 192 × 176, in-plane GRAPPA acceleration = 4. For P1-P15, high-resolution T1w and matched T2w scans were also acquired using Human Connectome Project (HCP) parameters. The T1w HCP ME-MPRAGE structural scan used: voxel size = 0.8 mm, TR = 2,500 ms, TE = 1.81, 3.60, 5.39, and 7.18 ms, TI = 1,000 ms, flip-angle = 8°, matrix 320 × 320 × 208, 144, in-plane GRAPPA acceleration = 2. The T2w HCP structural scan used a sampling perfection with application-optimized contrasts using different flip angle evolution sequence (SPACE, Siemens Healthineers, Erlangen, Germany): voxel size = 0.8 mm, TR=3,200 ms, TE=564 ms, 208 slices, matrix=320 × 300 × 208, in-plane GRAPPA acceleration = 2.

All resting-state fixation runs were screened for quality as described by Du and colleagues (2024).

### Data Processing and Registration that Minimizes Spatial Blurring

Data were processed using the openly available preprocessing pipeline (“iProc”) that preserved spatial details by minimizing spatial blurring and the number of interpolations (described in detail in Braga et al. 2019 and Du et al. 2024 with relevant methods repeated here). For S1 and S2, the pre-processed data were taken from Xue et al. (2021) and additionally processed using the 15-network cerebral network estimates reported in Du et al. (2024). For P1-P15, the pre-processed data and network estimates for the cerebral cortex were taken directly from Du et al. (2024).

Data were interpolated to a 1-mm isotropic native-space atlas (with all processing steps composed into a single interpolation) that was then projected using FreeSurfer v6.0.0 to the fsaverage6 cortical surface (40,962 vertices per hemisphere; (Fischl 2012; Fischl, Sereno, and Dale 1999)). Five transformation matrices were calculated: (1) a motion correction matrix for each volume to the run’s middle volume [linear registration, 6 degrees of freedom (DOF); MCFLIRT, FSL], (2) a matrix for field-map-unwarping the run’s middle volume, correcting for field inhomogeneities caused by susceptibility gradients (FUGUE, FSL), (3) a matrix for registering the field-map-unwarped middle BOLD volume to the within-individual mean BOLD template (12 DOF; FLIRT, FSL), (4) a matrix for registering the mean BOLD template to the participant’s T1w native-space image (6 DOF; using boundary-based registration, FreeSurfer), and (5) a non-linear transformation to MNI space (nonlinear registration; FNIRT, FSL). The individual-specific mean BOLD template was created by averaging all field-map-unwarped middle volumes after being registered to an up-sampled 1.2 mm and unwarped mid-volume template (interim target, selected from a low motion run acquired close to a field map). The T1w native-space template was then resampled to 1.0 mm isotropic resolution.

Confounding variables including 6 head motion parameters, whole-brain, ventricular signal, deep cerebral white matter signal, and their temporal derivatives were calculated from the BOLD data in T1w native space. The signals were regressed out from the BOLD data using 3dTproject, AFNI (Cox 1996, 2012). The residual BOLD data were then bandpass filtered at 0.01-0.10-Hz using 3dBandpass, AFNI (Cox 1996, 2012). For surface analyses, the native space data were resampled to the fsaverage6 standardized cortical surface mesh using trilinear interpolation and then surface-smoothed using a 2-mm full-width-at-half-maximum (FWHM) Gaussian kernel. For striatal analyses, we elected to use unsmoothed MNI volume space data due to signal blur at and around the VS (see Kosakowski et al. 2024). The iProc pipeline thus allowed for robustly aligned BOLD data with minimal smoothing. Relevant final output spaces included the MNI volume space and the fsaverage6 cortical surface.

### Striatum Identification and Visualization

As described in Kosakowski et al. (2024) and repeated here, a key step for the present inquiry was to ensure high-quality segmentations of the striatum within individual participants. To isolate striatal voxels, we identified the caudate, putamen, and NAc in each participant using the FreeSurfer automated parcellation (Fischl 2012). The FreeSurfer-based masks were combined, binarized, and warped to the space of the Montreal Neurological Institute (MNI; Evans et al. 1993) atlas using the same individual-specific analysis pipeline used in “iProc” preprocessing (Braga et al. 2019). Striatum masks were created for different purposes that included: (1) visualization of the striatal boundaries and (2) visualization of the cortex adjacent to the striatum. To accommodate the different uses of striatal masks, the boundary of the striatum was dilated at different levels using *fslmaths -dilM* and binarized. 1x dilation was used for striatal boundary visualization and 5x dilation was used to visualize the adjacent cerebral cortex.

### Within-Individual Striatum-to-Cortex Correlation Matrices

For each participant, the pair-wise Pearson correlation coefficients between the fMRI time courses at each surface vertex were calculated for each resting-state fixation run, yielding an 81,924 × 81,924 matrix (40,962 vertices / hemisphere). For subcortex, we used a volume mask that included striatum, thalamus, amygdala, and midbrain. For each fixation run, we computed pair-wise Pearson correlation coefficients for the fMRI time courses between each volume voxel within the mask and each cortical vertex. The subcortical matrix was 134,797 × 134,797 and the subcortical to cortical matrix was 134,797 × 81,924. The matrices were then Fisher *r*-to-*z* transformed and averaged across all runs to yield a single best estimate of the within-individual correlation matrices. These *z*-scored matrices were combined and assigned to a cortical and subcortical template combining left and right hemispheres of the fsaverage6 surface and subcortex of the MNI152 volume into the CIFTI format to interactively explore correlation maps using the Connectome Workbench’s wb_view software (Glasser et al. 2013; Marcus et al. 2011). The colorbar scales of correlation maps were thresholded at *z*(*r*)=0.07-0.20 using the Jet look-up table for visualization. Correlation maps are additionally provided on BALSA to enable viewing in alternative color scales.

### Individualized Network Estimates of the Cerebral Cortex

Striatal mapping began by utilizing within-individual precision maps of network organization in the cerebral cortex. For all participants, the 15-network cerebral cortex estimates were taken directly from Du et al. (2024) with relevant method description repeated here. A Multi-Session Hierarchical Bayesian Model (MS-HBM) was implemented to estimate the cortical networks (Kong et al. 2019). For S1 and S2, the resting-state fixation data were split into three subsets. For S1, Data Set 1 included runs 1-20, Data Set 2 included runs 21-40, and Data Set 3 included runs 41-62. For S2, Data Set 1 included runs 1-20, Data Set 2 included runs 21-40, and Data Set 3 included runs 41-61. The MS-HBM was independently implemented on each data subset. For P1-P15, the MS-HBM was implemented on each participant’s full set of resting-state fixation data.

To estimate networks in the cerebral cortex, the connectivity profile of each vertex on the cortical surface was first estimated as its functional connectivity to 1,175 regions of interest (ROIs) that uniformly distributed across the fsaverage6 surface meshes (Yeo et al. 2011). For each run of data, the Pearson’s correlation coefficients between the fMRI time series at each vertex (40,962 vertices / hemisphere) and the 1,175 ROIs were computed. The resulting 40,962 × 1,175 correlation matrix per hemisphere was then binarized by keeping the top 10% of the correlations to obtain the functional connectivity profiles.

Next, the MS-HBM was initialized with a group-level parcellation estimated from a subset of the HCP S900 data release using the clustering algorithm (Yeo et al. 2011). The group-level parcellation was used to initialize the expectation-maximization algorithm for estimating parameters in the MS-HBM. The goal of applying the model as used here was to obtain the best estimate of networks within each individual participant’s dataset, not to train parameters and apply them to unseen data from new participants (Kong et al. 2019).

The 15-network estimates include : Somatomotor-A (SMOT-A) network, Somatomotor-B (SMOT-B) network, Premotor-Posterior Parietal Rostral (PM-PPr) network, CG-OP network, SAL network, Dorsal Attention Network-A (dATN-A), Dorsal Attention Network-B (dATN-B), Frontoparietal Network-A (FPN-A), Frontoparietal Network-B (FPN-B), Default Network-A (DN-A), Default Network-B (DN-B), Language (LANG) network, Visual Central (VIS-C) network, Visual Peripheral (VIS-P) network, and Auditory (AUD) network. Here we focus on the SAL network and its relation to the CG-OP network. In several analyses, to provide context and insight into relations between adjacent networks, we also visualize and plot data from the FPN-A, FPN-B, LANG network, DN-B, DN-A, dATN-A, and dATN-B.

### Seed-Region Based Exploration of the Ventral Striatum

For S1 and S2 we identified a seed region in the VS in Data Set 1 that was maximally correlated with the SAL network. Then, we replicated the correlations of this *a priori* seed region in the replication Data Set 2 and triplication Data Set 3 from each participant. In P1-P15, we identified similar seed regions in the VS that were correlated with the SAL network using the full dataset from each individual.

For S1 and S2, in Data Set 1 we identified a second seed region in the ventral portion of the putamen (for brevity, we will refer to this region as the “ventral putamen”) that was maximally correlated with the CG-OP network. We replicated the correlations of this seed region in the replication Data Set 2 and triplication Data Set 3 for both participants. In P1-P15 we identified similar seed regions in the ventral putamen that were correlated with the CG-OP network using the full dataset from each individual.

### Control Analysis of Subgenual Cingulate

A key challenge of estimating networks in the VS is signal blur from the nearby cortical regions, particularly along the ventral midline. To check that signal blur was not a contributor, we performed a control analysis by which seed regions were placed in MPFC, specifically the region at or near the sgACC. For S1 and S2, the seed region was placed at MNI coordinates X = -4, Y = 16, Z = -8. We visualized the correlations at and near the striatum boundaries to determine if the correlation pattern extended into the VS. The surface correlations were also plotted to illustrate the full cortical pattern.

### Software and Statistical Analysis

Functional connectivity was calculated in MATLAB (version 2019a; MathWorks, Natick, MA) using Pearson’s product moment correlations. FreeSurfer v6.0.0, FSL, and AFNI were used during data processing. The estimates of networks on the cortical surface were visualized in Connectome Workbench v1.3.2. Model-free seed-region confirmations were performed in Connectome Workbench v1.3.2. Network parcellation was performed using code from (Kong et al. 2019) on Github. (https://github.com/ThomasYeoLab/CBIG/tree/master/stable_projects/brain_parcellation/Kong2019_MSHBM). All data are publicly available on NIMH Data Archive (NDA) and striatal results are available on BALSA; all code and data for generating Figures is available on Open Science Framework (OSF).

## Results

### The Ventral Striatum Correlates with Distributed Cortical Regions that Recapitulate the Full Extent of the Salience Network

The main result of this paper is that the VS is coupled to a distributed network of regions in the cerebral cortex that is robustly and preferentially aligned to the SAL network. This result is revealed in multiple ways across all participants examined.

In the initial two participants (S1 and S2), seed regions in the VS were manually identified in the first datasets (Data Set 1) that were strongly correlated with the SAL network (Figures 1 and 2). The distributed network prominently included regions of the SAL network in the anterior insula (indicated with an asterisk in Figures 1 and 2), the middle and anterior cingulate, as well as posteromedial regions. In both participants the VS seed region was also strongly correlated with a region in DLPFC (indicated with an arrow in Figures 1 and 2) and a rostral region of the medial prefrontal cortex (MPFC; indicated with a double asterisk in Figures 1 and 2). Collectively these regions, including those in DLPFC, mostly fell within and near to the boundaries of the SAL network, including across the network’s entire distributed extent. There were also weak and non-specific correlations, including regions at or near retinotopic visual cortex.

**Figure 2.**
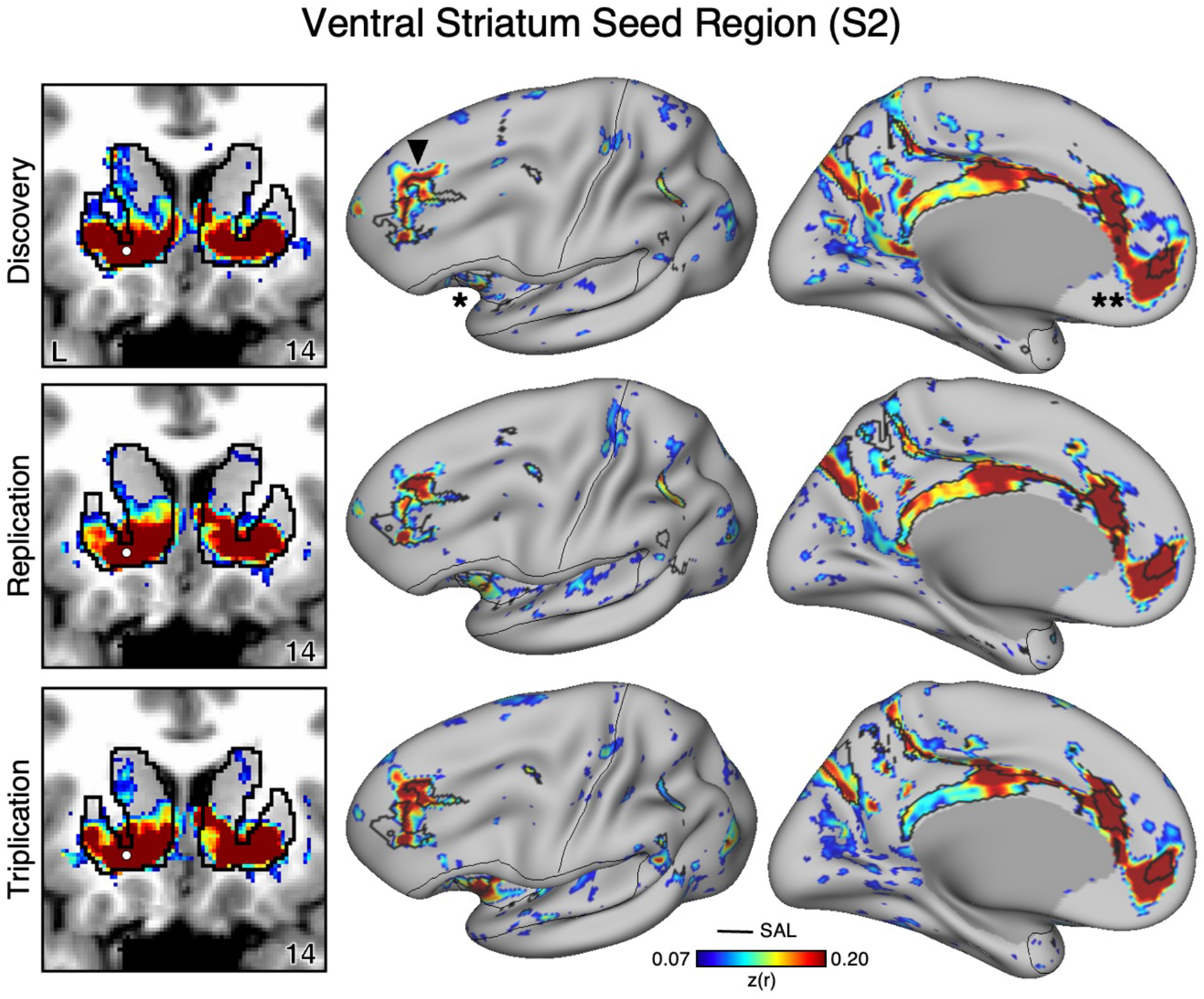
The Ventral Striatum (VS) Correlates with the Full Cortical Extent of the Salience (SAL) Network in S2. A seed region (indicated with a white circle) in the VS (top left) recapitulates the full extent of the SAL network, similar to S1 but with idiosyncratic spatial differences. In independent data from S2, this pattern was replicated (second row) and triplicated (third row) using the same seed region (white circles). Results are visualized and labeled as described in Figure 1.

Critically, the distributed pattern of correlation with the SAL network was independently replicated using the same seed region in Data Set 2 (2^nd^ row in Figures 1 and 2) and triplicated in Data Set 3 (3^rd^ row in Figures 1 and 2) in both participants. A striking correspondence was observed between the topography produced by the VS seed regions and the SAL network borders. In the volume, as a control, the correlations with the seed regions were examined and did not expand beyond striatal boundaries (Figures 1 and 2, left column) indicating the observed signal was unlikely due to bleed from nearby cortical structures (see Kosakowski et al. 2024). This was similar for all three datasets within each participant. Thus, while the seed regions were identified manually in the initial discovery Data Set 1, the detailed and selective patterns of correlations were found in multiple independent datasets from the same individuals.

### An Anatomically Adjacent Seed Region in the Ventral Portion of the Putamen Recapitulates the Full Extent of the Cingulo-Opercular Network

To assess the specificity of the VS seed regions that were correlated with the SAL network, we compared the topography produced by a seed region placed in an adjacent anatomical location - the ventral portion of the putamen, which we will refer to as “ventral putamen” for simplicity (for context, this region may fall within a homolog of VSd of the macaque monkey as described by Choi et al. 2017). In S1 and S2, we identified a seed region in the ventral putamen using discovery Data Set 1 that was maximally correlated with the CG-OP network (Figures 4 and 5, top row). The selected seed region reproduced the entire extent of the CG-OP network including canonical regions in the insula (indicated with an asterisks in Figures 4 and 5), cingulate cortex (indicated with double asterisks in Figures 4 and 5), recently discovered inter-effector regions (Gordon et al. 2023), as well as a region in DLPFC (indicated with an arrow in Figures 4 and 5). Importantly, this spatial pattern of correlation was replicated (2^nd^ row, Figure 3 and 4) and triplicated (3^rd^ row, Figure 3 and 4) in both participants using the same seed region applied to independent datasets (Data Sets 2 and 3). Correlations did not appear to expand beyond the boundaries of the striatum and avoided the region of VS associated with the SAL network from the previous analyses. These details suggest a dissociation between the functional connectivity of these adjacent striatal regions.

**Figure 3.**
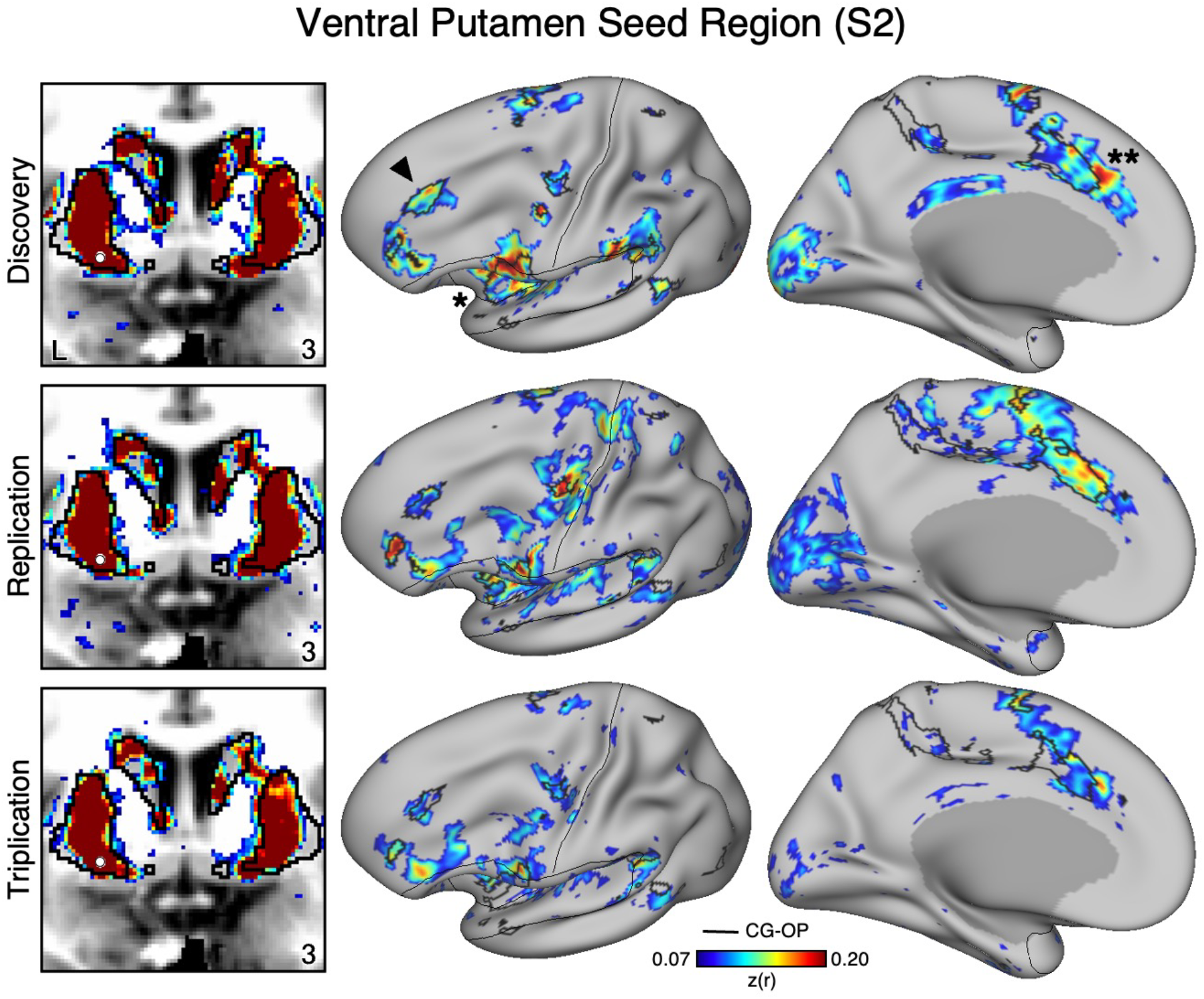
The Ventral Putamen Correlates with the Full Cortical Extent of the Cingulo-Opercular (CG-OP) Network in S1. A seed region (indicated with a white circle) in the ventral portion of the putamen (top left) recapitulates the full extent of the CG-OP network, including the DLPFC (indicated with a black arrow), insula (indicated by asterisk), and an anterior midline region near to premotor cortex (indicated by double asterisks). The pattern also includes multiple discontinuous regions along the pre-central gyrus (“inter-effector” regions). In independent data from S1, this pattern was replicated (second row) and triplicated (third row) using the same seed region (white circles). Cortical results are visualized on the inflated fsaverage6 surface with the lateral view slightly tilted to show the dorsal surface. Striatal results are visualized in coronal sections on each individual’s own T1-weighted anatomical image transformed to the atlas space of the MNI atlas. L = left. Coordinates in the bottom right of the left panels indicate the y coordinates (in mm). Black outlines in the volume indicate the boundaries of the striatum and outlines on the cortical surface indicate the CG-OP network (see methods). All correlation values are plotted after r to z conversion using the jet colorscale shown by the legend.

**Figure 4.**
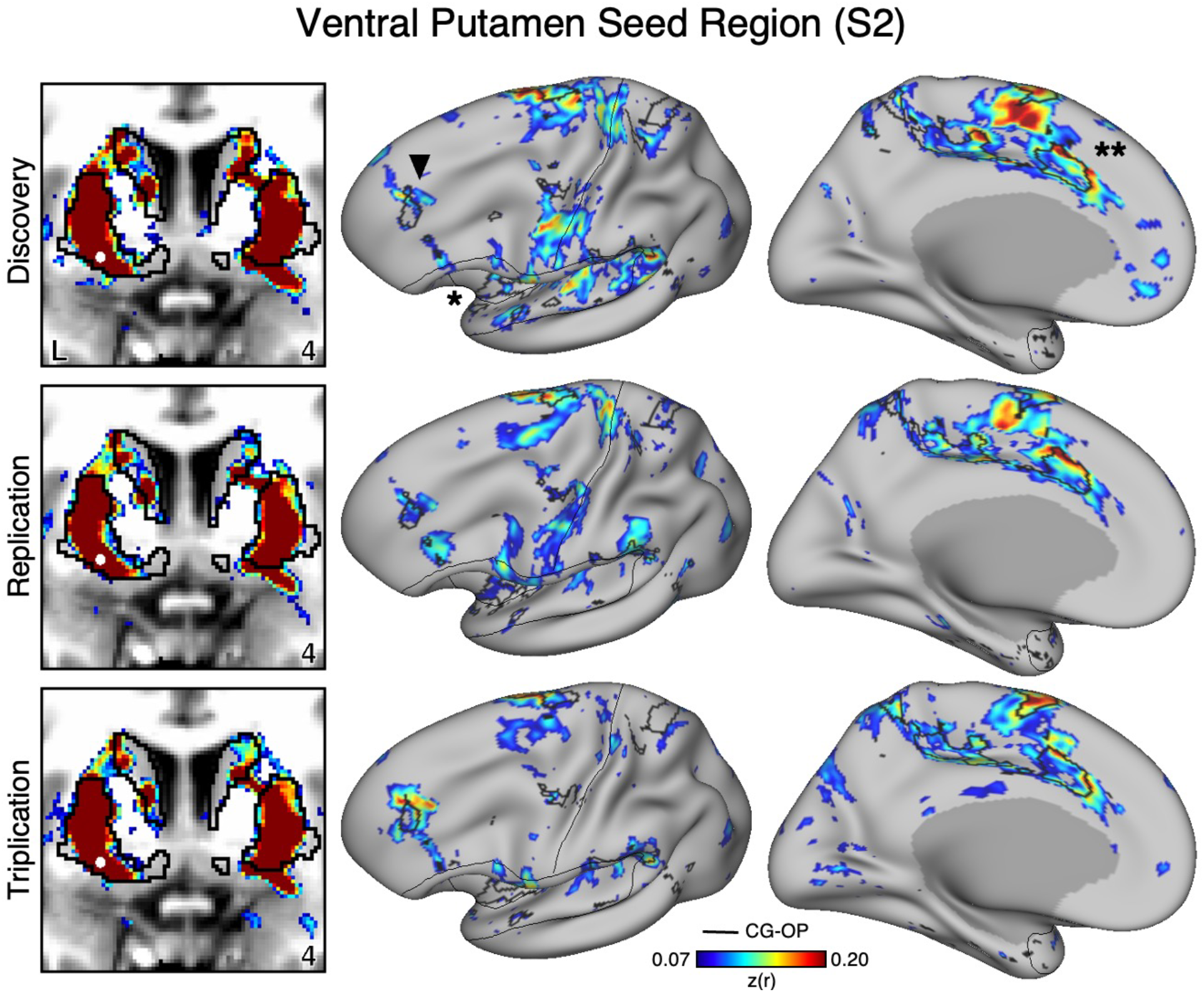
The Ventral Putamen Correlates with the Full Extent of the Cingulo-Opercular (CG-OP) Network in S2. A seed region (indicated with a white circle) in the ventral putamen (top left) recapitulates the full extent of the CG-OP network, similar to S1 but with idiosyncratic spatial differences. In independent data from S2, this pattern was replicated (second row) and triplicated (third row) using the same see region (white circles). Results are visualized and labeled as described in Figure 3.

**Figure 5.**
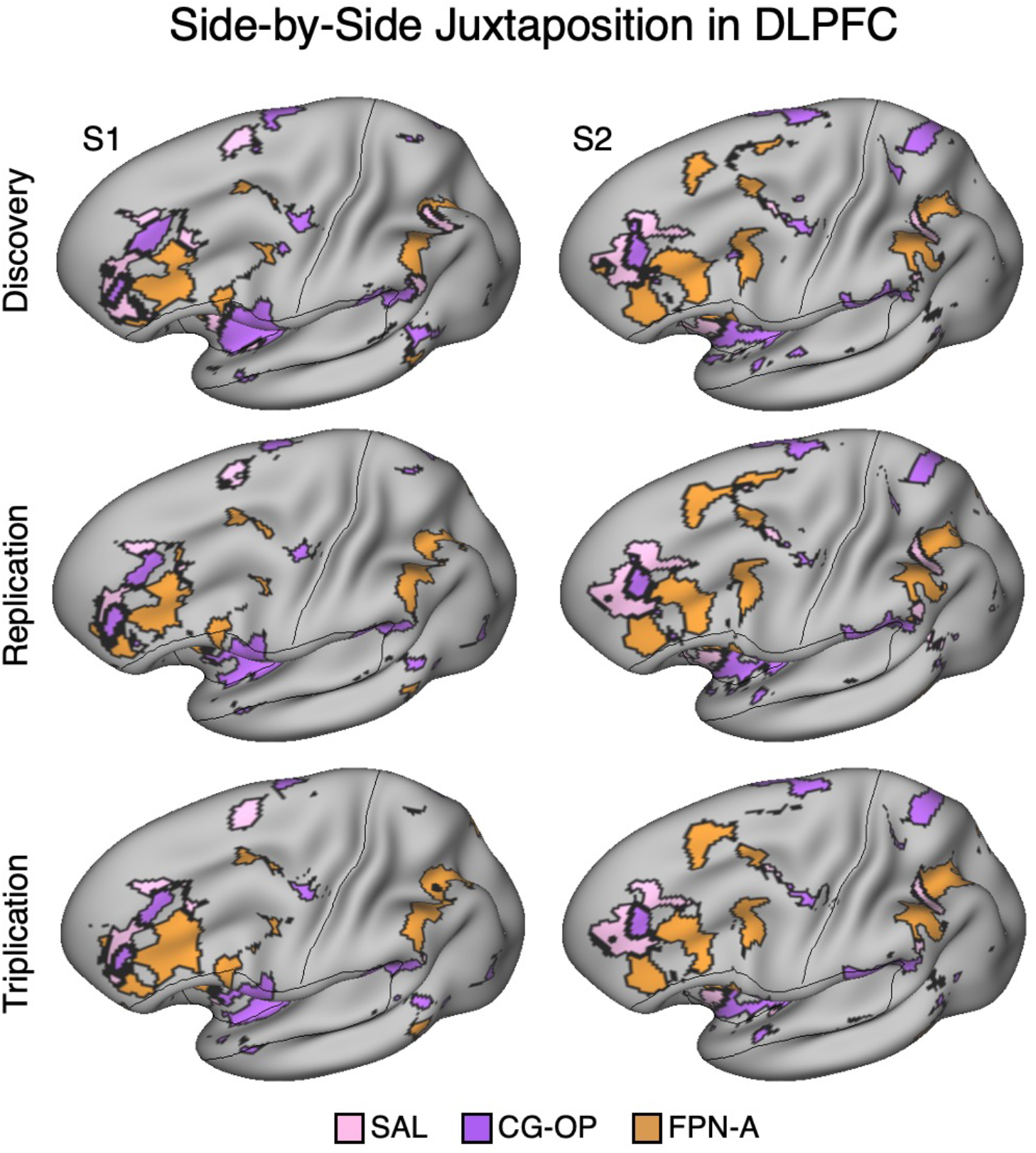
Side-by-Side Adjacency of Distinct Network Regions in the Dorsolateral Prefrontal Cortex (DLPFC) of S1 and S2. In each Data Set from S1 and S2, the SAL network (pink), CG-OP network (purple), and FPN-A (orange) have distinct representations in DLPFC that are juxtaposed with one another. The DLPFC portions of the SAL and CG-OP networks are rostral to the FPN-A regions. Cortical results are visualized on the inflated fsaverage6 surface slightly tilted to show the dorsal surface. All networks are bilateral but only the left hemisphere surface is displayed.

### Salience, Cingulo-Opercular, and Frontoparietal Networks are Adjacent in the DLPFC

In three independent datasets from S1 and S2, a seed region in the VS was positively correlated with the SAL network, including a specific region in DLPFC. Additionally, a nearby seed region in the ventral putamen was positively correlated with the CG-OP network, including a distinct region in DLPFC. The DLPFC is often associated in the human literature with Frontoparietal Network-A (FPN-A)^l^, a network that supports domain flexible cognitive control (Braga et al. 2020; Du et al. 2024; Duncan 2010; Fedorenko, Duncan, and Kanwisher 2013). Thus, we next visualized the MS-HBM defined boundaries of the SAL network, CG-OP network, and FPN-A in DLPFC. Regions associated with the SAL network, CG-OP network, and FPN-A are tightly juxtaposed in DLPFC in all three independent datasets from both participants (Figure 5). In each participant, the SAL network is linked to a more rostral region, FPN-A is linked to a more caudal region, and the region linked to the CG-OP network is found roughly between the SAL network and FPN-A, although the topography is complex and idiosyncratic in each of the two individuals.

### Ventral Striatum and Ventral Putamen Seed Regions Correlate with Adjacent DLPFC Regions

How specific is the correlation between VS and the SAL network regions in DLPFC? As a secondary visualization of the results plotted in Figures 1 and 2, the mean correlations from the VS seed region are displayed for multiple networks -first within only the DLPFC and then across the full distributed extent of each network. The DLPFC regional correlations with the VS seed region from Data Set 2 and Data Set 3 were the greatest for the SAL network region (Figure 6, top left). There was also a strong positive correlation with the DLPFC region linked to the CG-OP network. Interestingly, despite the tight juxtaposition of the SAL network and FPN-A in DLPFC, the VS seed region had a negative correlation with the FPN-A region in DLPFC.

**Figure 6.**
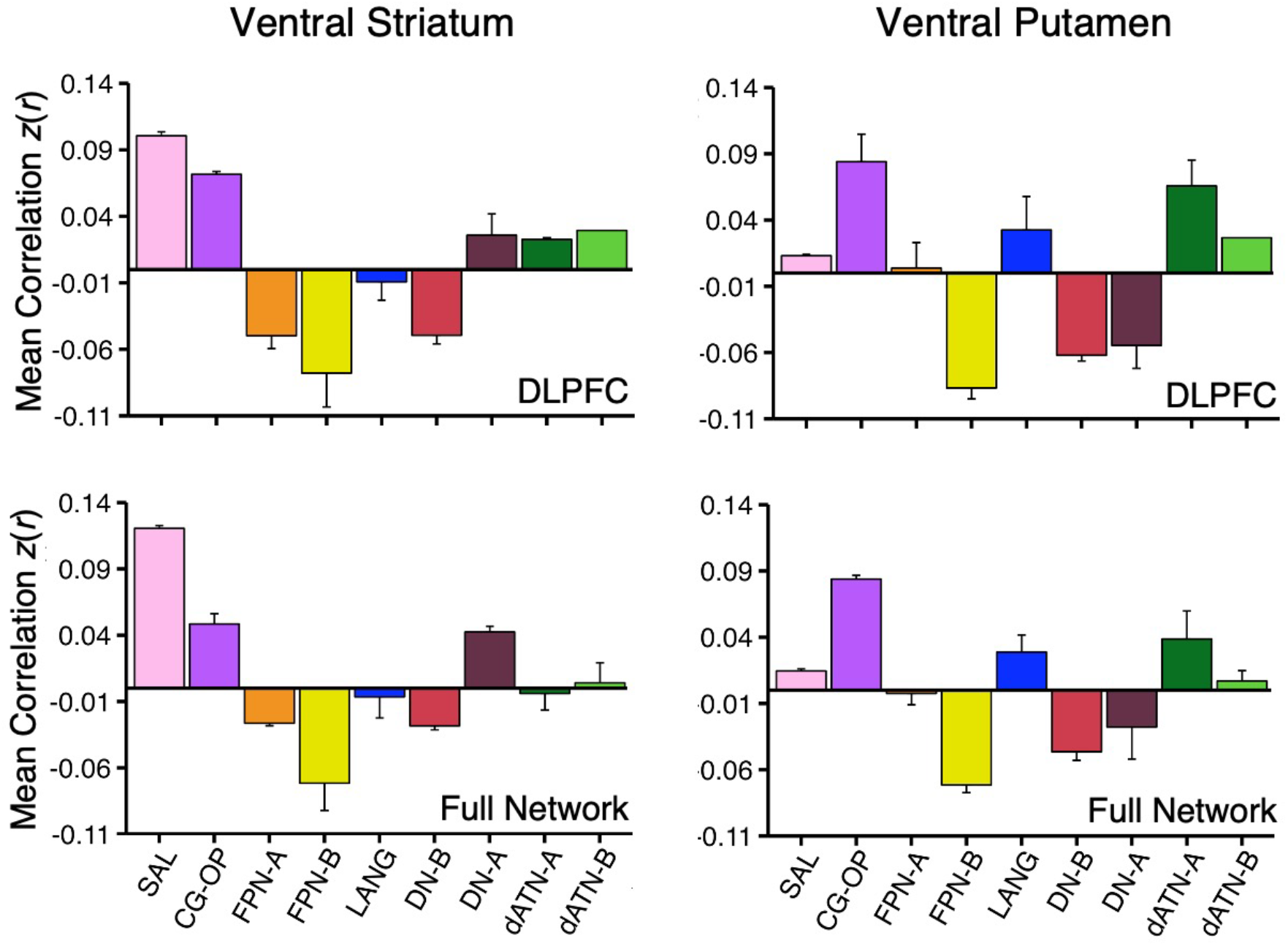
Ventral Striatum (VS) and Ventral Putamen Seed Regions Preferentially Correlate with Distinct Networks. Mean correlation values are plotted for the DLPFC (Top Panel) and full cortical networks (Bottom Panel) from the two separate striatal seed regions (Left Column, VS; Right Column, Ventral Putamen). The correlation values are the mean values from S1 and S2 extracted from Data Sets 2 and 3 which were independent from the data used to select the seed regions. The VS seed regions are maximally correlated with the SAL network, while the ventral putamen seed regions are maximally correlated with the CG-OP network. The VS seed regions also show strong, positive correlation with the CG-OP network showing the pattern is preferential not selective. Bars show mean correlation (*z*(*r*)) in each network, error bars indicate standard error of the mean adjusted for within-subject variance (Cousineau 2005).

The VS seed region was also negatively correlated with FPN-B and DN-B regions in DLPFC. This pattern of positive and negative correlations was recapitulated in the full extent of the cerebral cortex (Figure 6, bottom left). Thus, the VS is linked to specific regions preferentially associated with the SAL and CG-OP networks, and not networks associated with cognitive control (e.g., FPN-A).

The ventral putamen seed region had the strongest correlation with the CG-OP network region in DLPFC (Figure 6, top right). The ventral putamen seed region also had strong positive correlations with dATN-A and LANG network regions in DLPFC and negative correlations with FPN-B, DN-B, and DN-A regions in DLPFC. Despite the tight juxtaposition between DLPFC regions of the CG-OP network and FPN-A, the ventral putamen seed region had a near zero correlation with the FPN-A region. This was also the case in the full cortical extent of each network (Figure 6, bottom right).

Thus, while the VS and ventral putamen seed regions were both associated with the SAL and CG-OP networks, which possess tightly juxtaposed regions in DLPFC, there was a reliable difference such that the VS was more correlated with the SAL network than the CG-OP network, and the ventral putamen was more correlated with the CG-OP network than the SAL network. Both the VS and ventral putamen were absent positive correlations with DLPFC regions linked to FPN-A.

### The Subgenual Anterior Cingulate has Minimal Correlations with the Ventral Striatum

The VS and ventral putamen seed regions are preferentially correlated with the SAL and CG-OP networks. Neither seed region displayed correlations with ventral MPFC including the region at or around sgACC. To better understand the relation between sgACC and VS, we visualized correlations with the sgACC seed region in each dataset. In both participants (S1 and S2), the sgACC seed region was robustly and preferentially correlated with DN-A but correlations with the VS were minimal and localized to the ventral boundary of the VS (Figures 7-8). This pattern of correlations was replicated in all three datasets for both participants. Along with our previous findings (Kosakowski et al. 2024), these results suggest that a strong interpretation of links between DN-A and VS is beyond the resolution of our data and that previous results, especially at the group level, may have been corrupted by signal blur from the nearby cortex. The present within-individual analysis approach appears to mitigate signal blur from MPFC into the VS.

**Figure 7.**
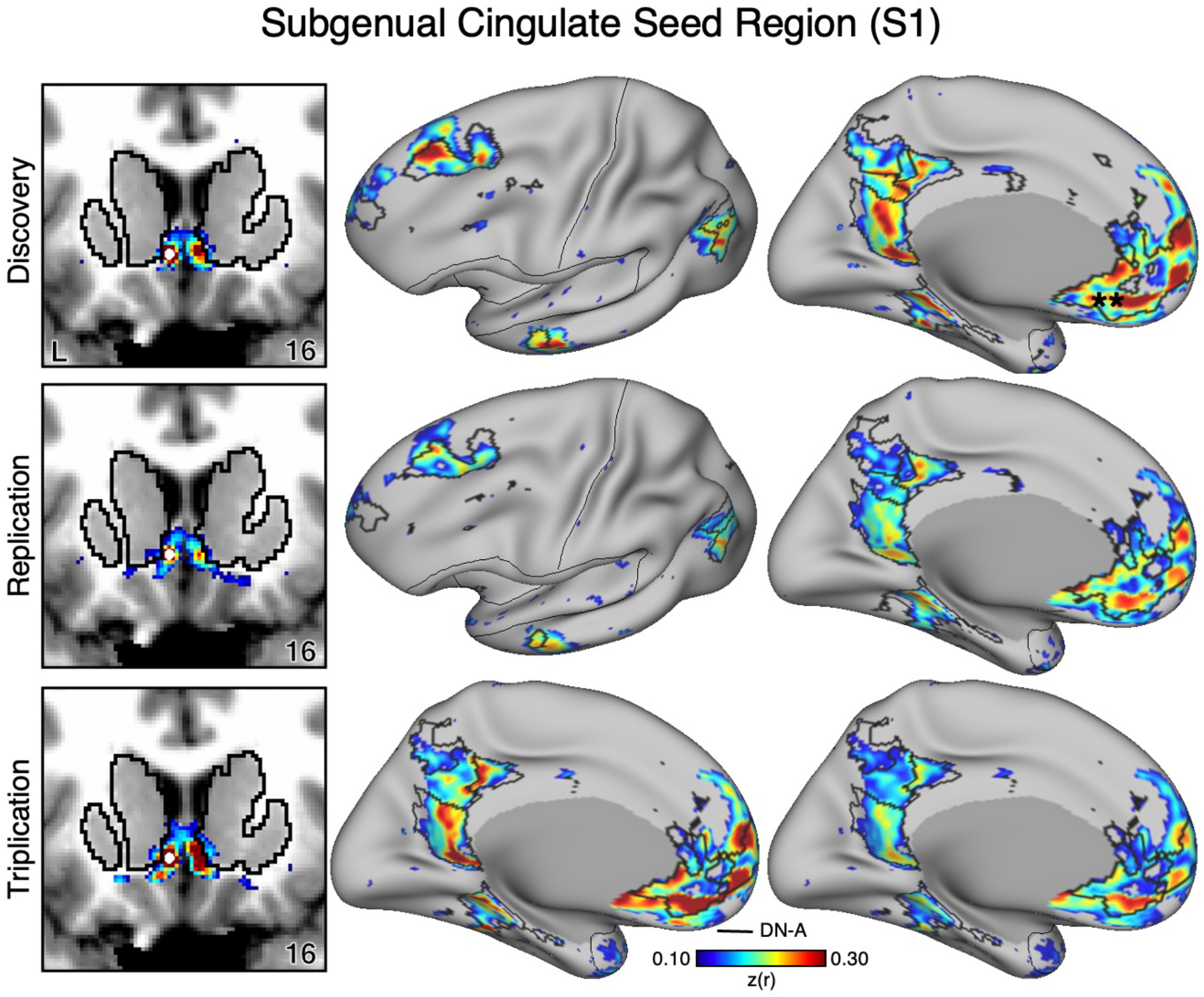
The Subgenual Anterior Cingulate (sgACC) has Minimal Correlations with the Ventral Striatum (VS) in S1. The VS is adjacent to the MPFC including the sgACC. As a control analysis, the correlations from an sgACC seed region (indicated with a white circle) were visualized to examine whether the correlation pattern extended into the VS (which is observed in group-averaged data) in S1. The sgACC seed region is strongly correlated with the full cortical extent of Default Network-A (DN-A), including regions in DLPFC (middle and right). Critically, the correlation pattern from sgACC minimally crosses the striatal boundary into the ventral portion of the VS (left). Cortical results are visualized on the inflated fsaverage6 surface with the lateral view slightly tilted to show the dorsal surface. Striatal results are visualized on each individual’s own T1-weighted anatomical image transformed to the atlas space of the MNI atlas. L = left. Coordinates in the bottom right of the left panels indicate the y coordinates (in mm). Black outlines in the volume indicate the boundaries of the striatum and outlines on the cortical surface indicate DN-A (see methods). All correlation values are plotted after r to z conversion using the jet colorscale shown by the legend.

**Figure 8.**
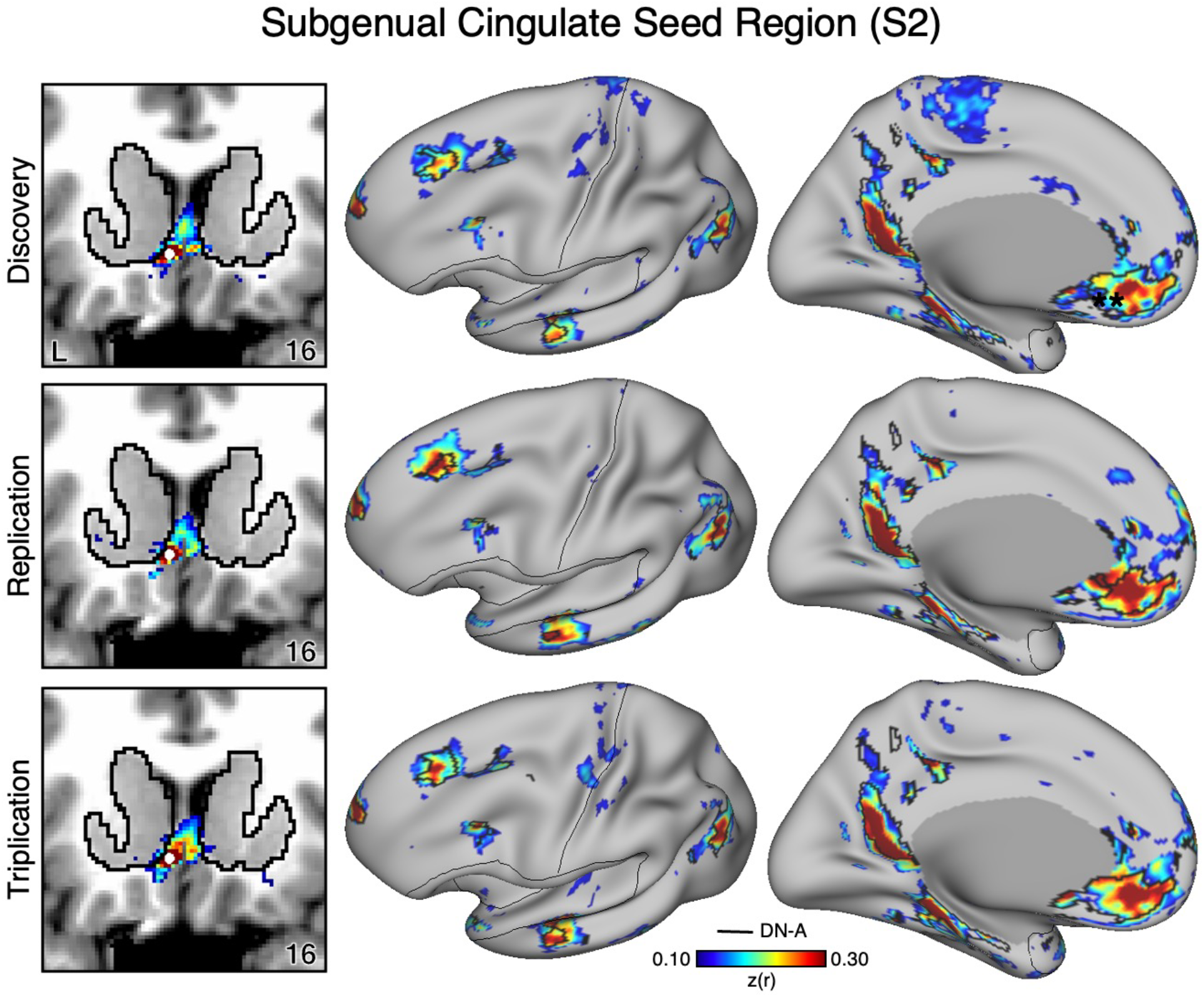
The Subgenual Anterior Cingulate (sgACC) has Minimal Correlations with the Ventral Striatum (VS) in S2. As a control analysis, the correlations from an sgACC seed region (indicated with a white circle) were visualized to examine whether the correlation pattern extended into the VS (which is observed in group-averaged data) in S2. The sgACC seed region is again strongly correlated with the full cortical extent of Default Network-A (DN-A; right) and minimally crosses the striatal boundary into the ventral portion of the VS (left). Results are visualized and labeled as described in Figure 7.

### Correlations Between the VS and the Salience Network Generalize to New Participants

To determine if VS correlations with the SAL network generalize, we identified a seed region in the VS for 15 additional participants in the same manner as for the first two participants (Figures 9-11). In every new participant, a seed region in VS again recapitulated the full extent of the SAL network, including regions in DLPFC, posteromedial cortex, anterior insula, and rostral MPFC. The selected VS seed regions had little or no observable correlations that extended beyond striatal boundaries into the cortex indicating observed signal likely originated from the VS and not adjacent cortex (Figures 9-11, left column). Thus, while the analysis did not allow for within-individual replication, each new participant analyzed in the same manner as the first two individuals possessed a VS region that was strongly correlated with the full extent of the SAL network.

**Figure 9.**
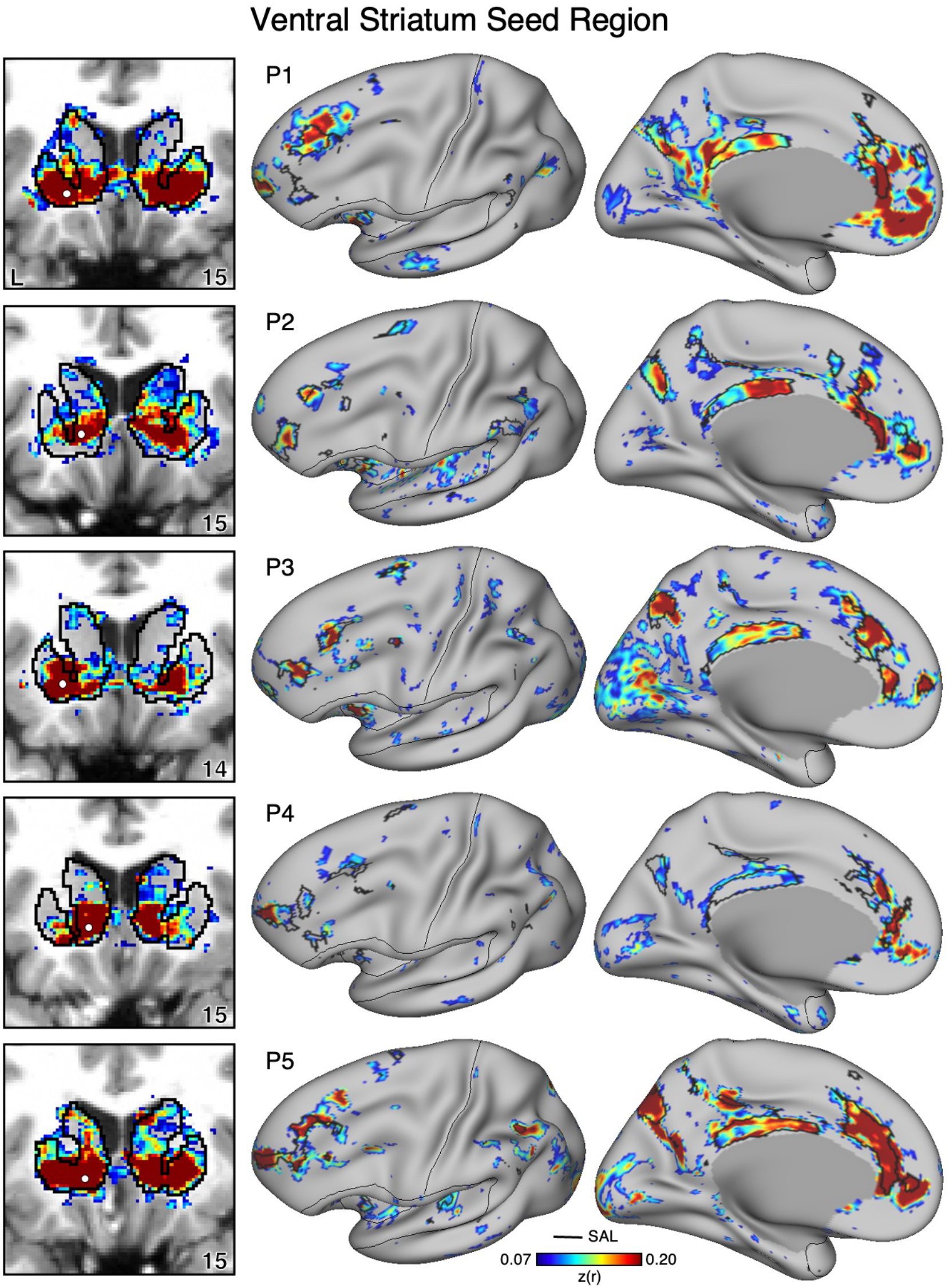
Ventral Striatum (VS) Correlates with the Full Cortical Extent of the Salience (SAL) Network in P1-P5. In new participants (P1-P5), seed regions (indicated with white circles) in the VS (left) recapitulate the full extents of the SAL network (right). Cortical results are visualized on the inflated fsaverage6 surface with the lateral view slightly tilted to show the dorsal surface. Striatal results are visualized on each individual’s own T1-weighted anatomical image transformed to the atlas space of the MNI atlas. L = left. Coordinates in the bottom right of the left panels indicate the y coordinates (in mm). Black outlines in the volume indicate the boundaries of the striatum and outlines on the cortical surface indicate the SAL network (see methods). All correlation values are plotted after r to z conversion using the jet colorscale shown by the legend.

**Figure 10.**
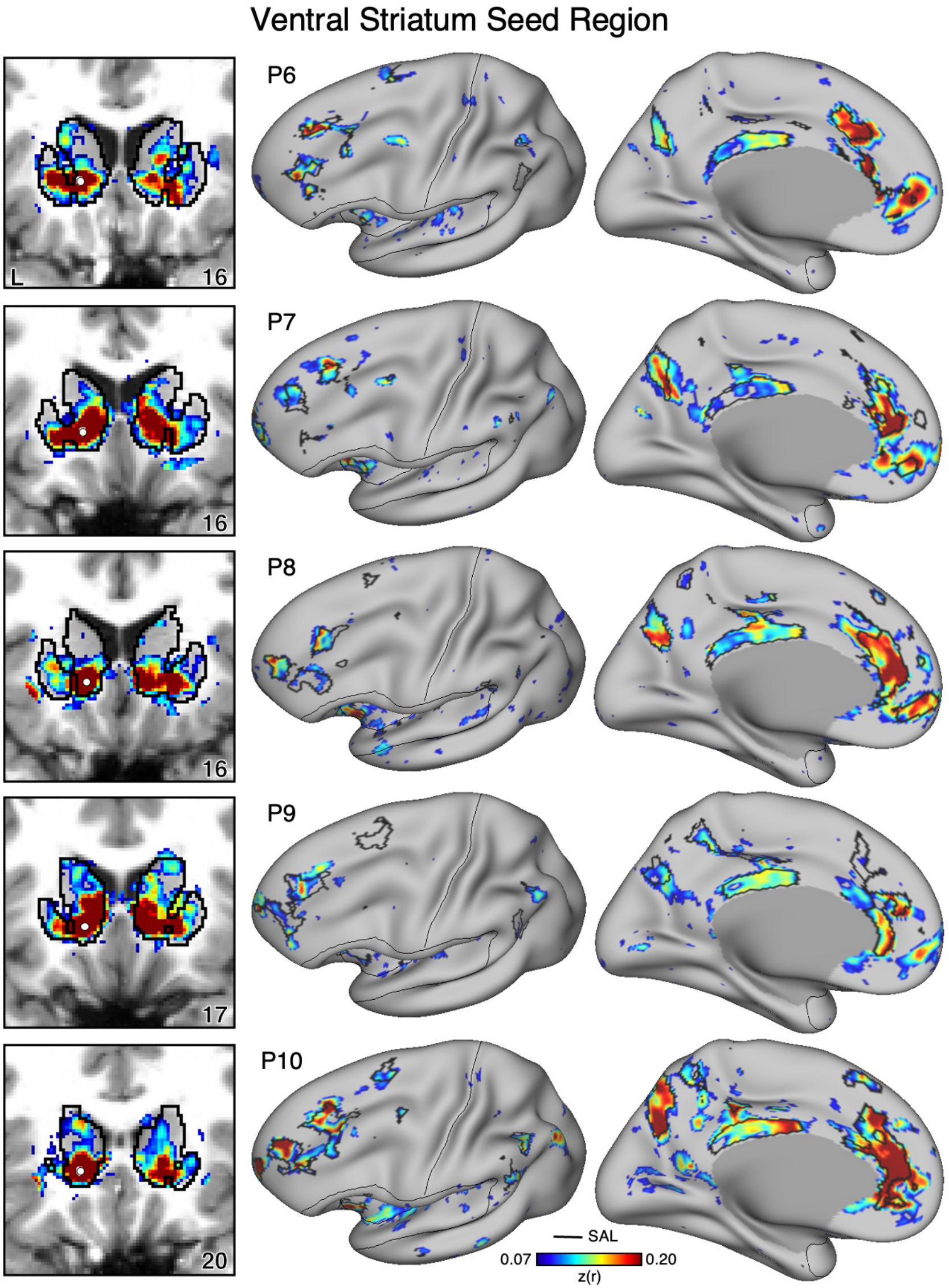
Ventral Striatum (VS) Correlates with the Full Cortical Extent of the Salience (SAL) Network in P6-P10. In additional new participants (P6-P10), seed regions (indicated with white circles) in the VS (left) again recapitulate the full extents of the SAL network (right). Results are visualized and labeled as described in Figure 9.

**Figure 11.**
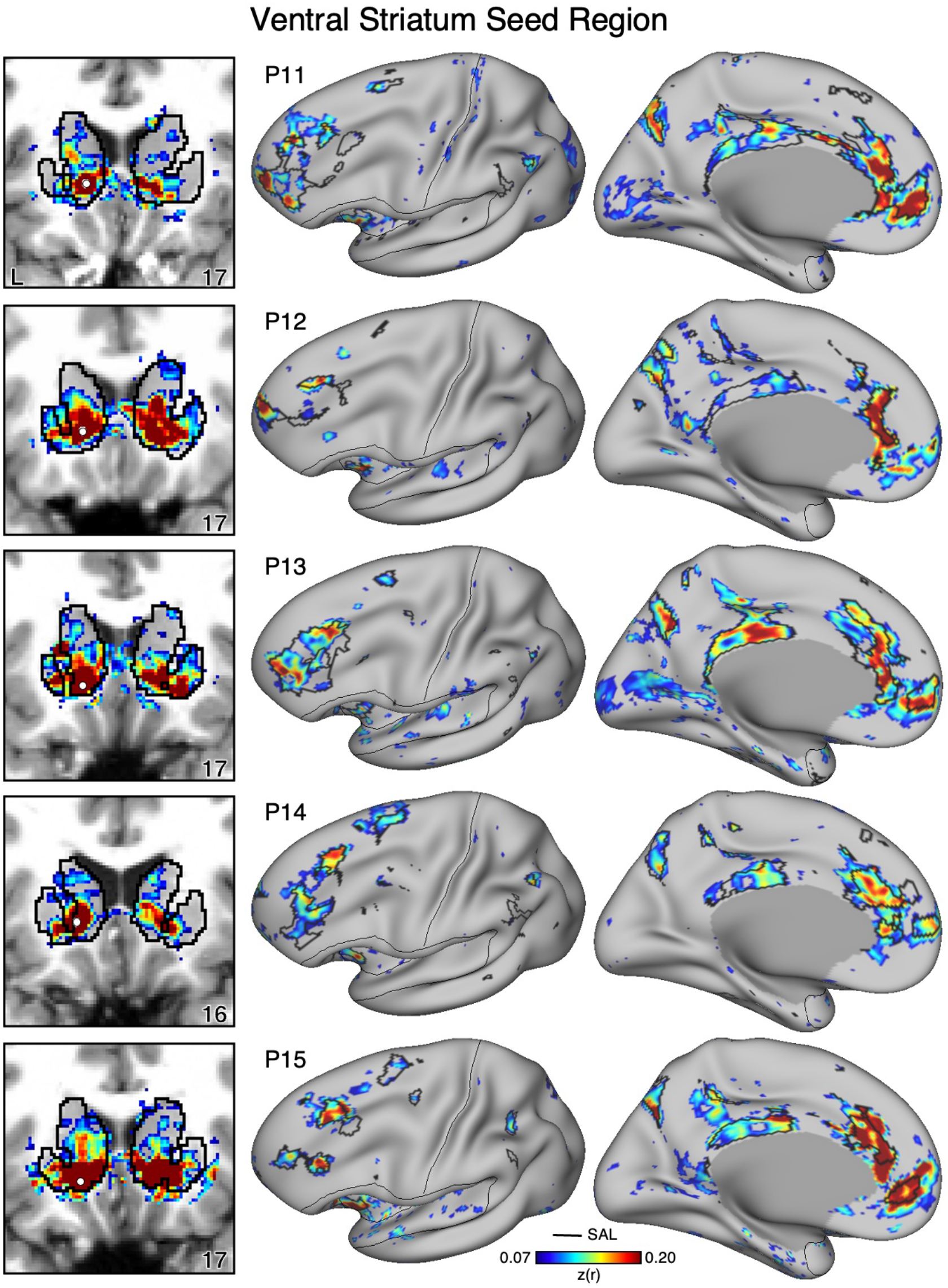
Ventral Striatum (VS) Correlates with the Full Cortical Extent of the Salience (SAL) Network in P11-P15. In a final set of new participants (P11-P15), seed regions (indicated with white circles) in the VS (left) again recapitulate the full extents of the SAL network (right). Results are visualized and labeled as described in Figure 9.

### Ventral Putamen Correlations with the Cingulo-Opercular Network Generalize to New Participants

To determine if ventral putamen correlations with the CG-OP network generalize, we repeated the same procedure as above for the ventral putamen. In every new participant, the ventral putamen seed region recapitulated the full extent of the CG-OP network (Figures 12-14), including distinct regions in DLPFC, insula, cingulate cortex, and the recently discovered inter-effector regions (Gordon et al. 2023).

**Figure 12.**
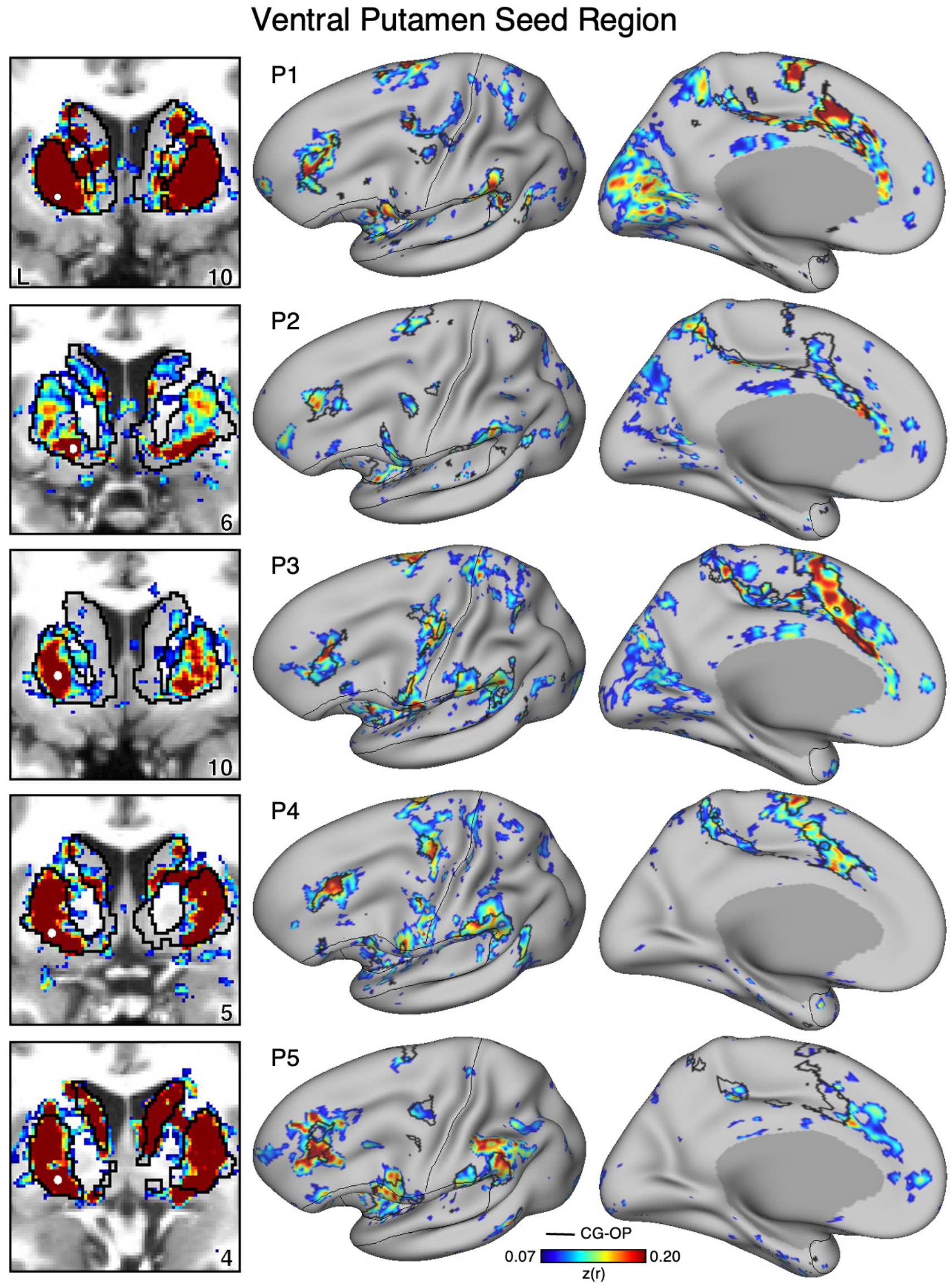
The Ventral Putamen Correlates with the Full Cortical Extent of the Cingulo-Opercular (CG-OP) Network in P1-P5. In new participants (P1-P5), seed regions (indicated with white circles) in the ventral portion of the putamen (left) recapitulate the full extents of the CG-OP network (right). Cortical results are visualized on the inflated fsaverage6 surface with the lateral view slightly tilted to show the dorsal surface. Striatal results are visualized on each individual’s own T1-weighted anatomical image transformed to the atlas space of the MNI atlas. L = left. Coordinates in the bottom right of the left panels indicate the y coordinates (in mm). Black outlines in the volume indicate the boundaries of the striatum and outlines on the cortical surface indicate the CG-OP network (see methods). All correlation values are plotted after r to z conversion using the jet colorscale shown by the legend.

**Figure 13.**
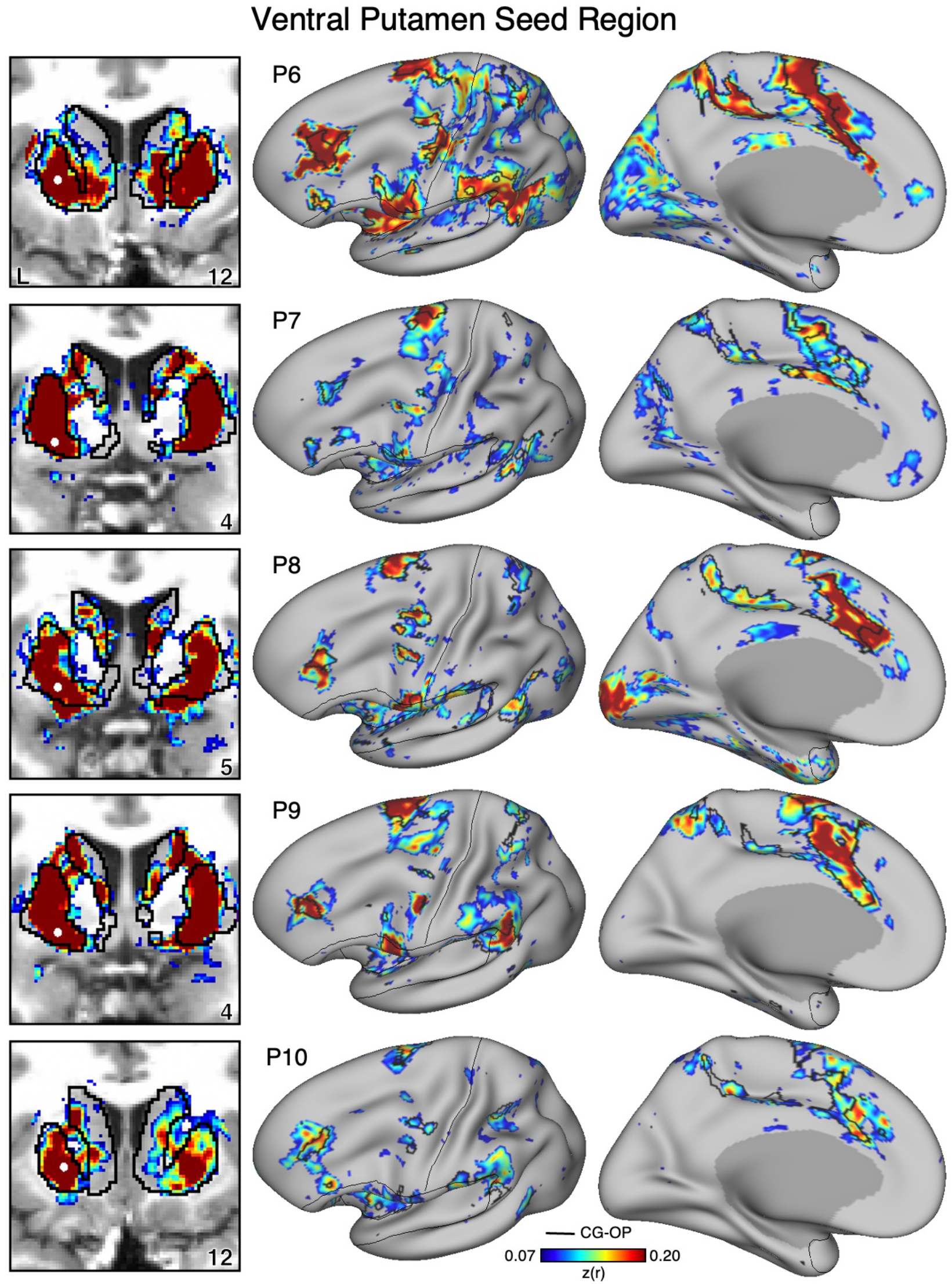
The Ventral Putamen Correlates with the Full Cortical Extent of the Cingulo-Opercular (CG-OP) Network in P6-P10. In additional new participants (P6-P10), seed regions (indicated with white circles) in the ventral portion of the putamen (left) again recapitulate the full extents of the CG-OP network (right). Results are visualized and labeled as described in Figure 12.

### Adjacency of the Salience, Cingulo-Opercular, and Frontoparietal Networks in the DLPFC Generalizes to New Participants

In each of the additional 15 participants, there are regions associated with the SAL network, the CG-OP network, and FPN-A in DLPFC. To again visualize the complex topography of the DLPFC, Figure 15 displays a rotated view of the network regions to fully visualize the DLPFC in the new participants. Despite idiosyncratic differences between participants, the juxtapositions of network regions in DLPFC showed generally similar spatial relations. The DLPFC region associated with the SAL network tended to be the most rostral, the FPN-A region more caudal, and the CG-OP network region nestled between them. Thus, there appears to be consistent general relations between network regions despite marked individual differences in the exact extents and positioning of the regions.

**Figure 14.**
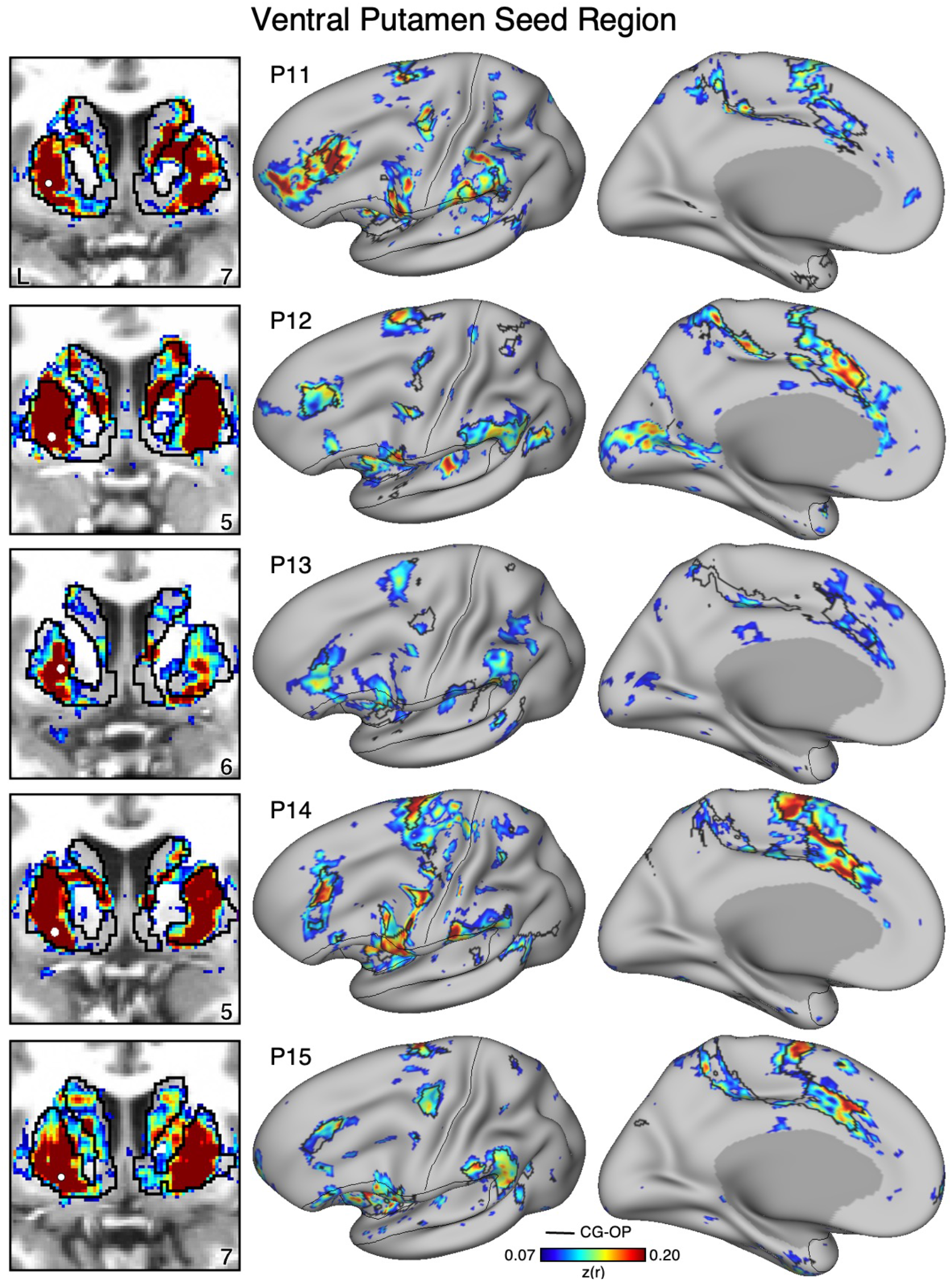
The Ventral Putamen Correlates with the Full Cortical Extent of the Cingulo-Opercular (CG-OP) Network in P11-P15. In additional new participants (P11-P15), seed regions (indicated with white circles) in the ventral portion of the putamen (left) again recapitulate the full extents of the CG-OP network (right). Results are visualized and labeled as described in Figure 12.

**Figure 15.**
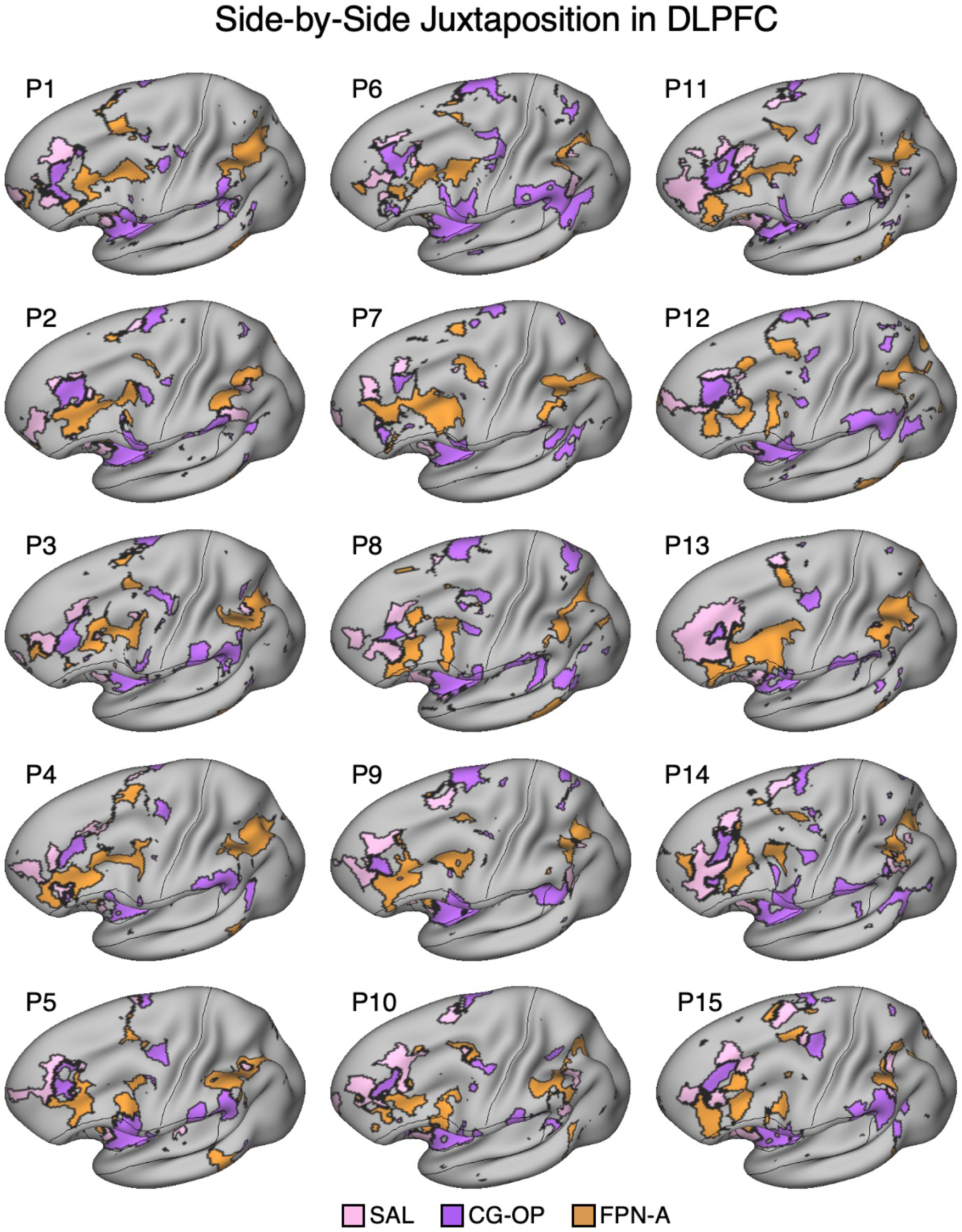
Side-by-Side Adjacency of Distinct Network Regions in the Dorsolateral Prefrontal Cortex (DLPFC) of P1-P15. In each participant (P1-P15), the SAL network (pink), the CG-OP network (purple), and FPN-A (orange) have distinct representations in DLPFC that are juxtaposed with one another. The DLPFC portions of the SAL and CG-OP networks are again rostral to the FPN-A regions. Cortical results are visualized on the inflated fsaverage6 surface slightly tilted to show the dorsal surface. All networks are bilateral but only the left hemisphere surface is displayed.

### Ventral Striatum and Ventral Putamen Seed Regions Correlate with Adjacent DLPFC Regions in New Participants

On average across all 15 participants, the VS seed region had the strongest correlation with the SAL network in DLPFC (Figure 16, top left) and across the full extent of the cerebral cortex (Figure 16, bottom left); the strength of correlations with the SAL network was similar in DLPFC alone and in the full extent of the SAL network. On average, the VS seed region was negatively correlated with the closely juxtaposed FPN-A region in DLPFC. Further, the VS seed region had a positive correlation with DN-A and a negative correlation with dATN-A.

**Figure 16.**
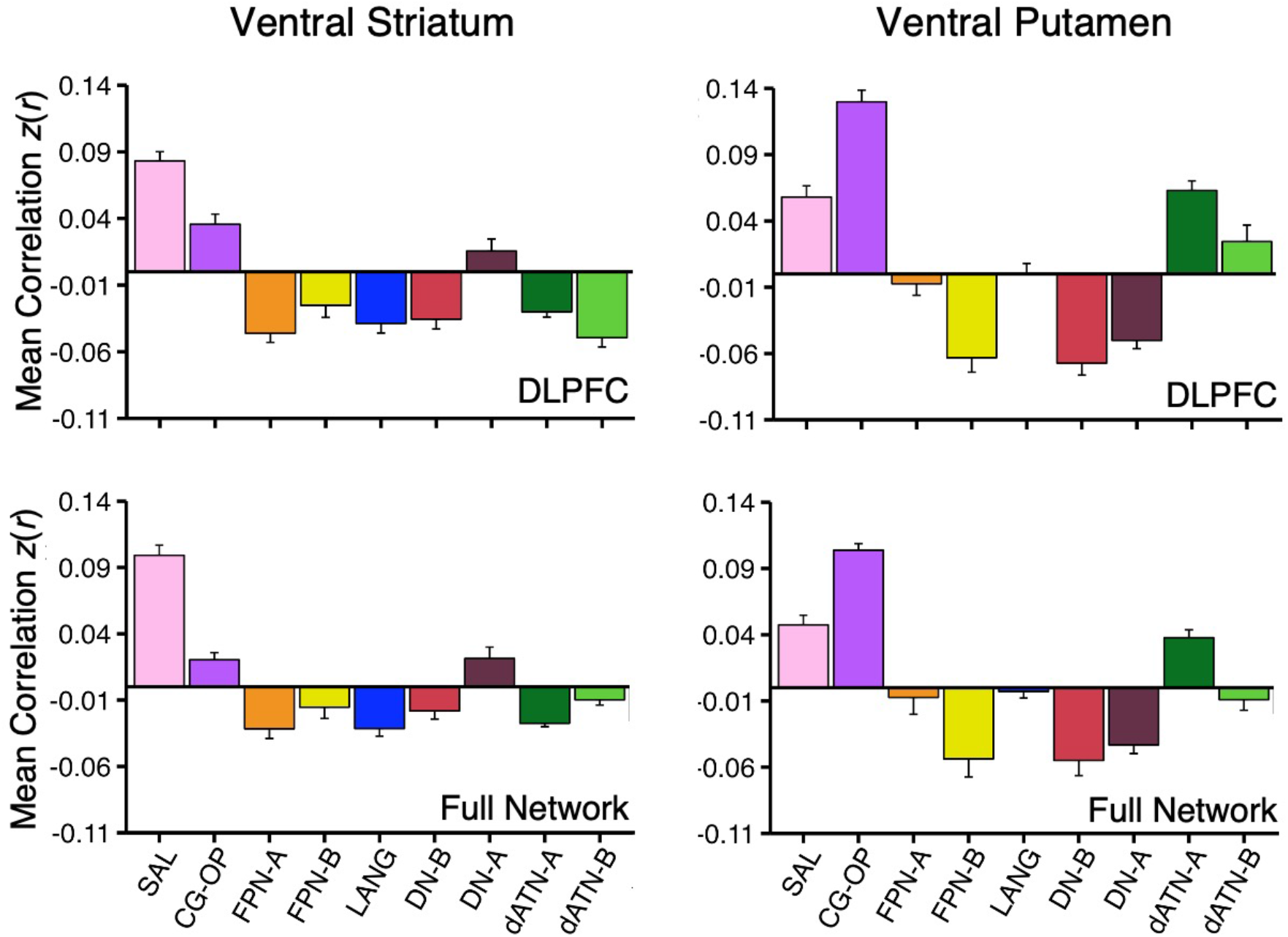
Ventral Striatum (VS) and Ventral Putamen Seed Regions Preferentially Correlate with Distinct Networks in P1-P15. Mean correlation values are plotted for the DLPFC (Top Panel) and full cortical networks (Bottom Panel) from the two separate striatal seed regions for all new participants (Left Column, VS; Right Column, Ventral Putamen). The correlation values are the mean values from P1-P15. The VS seed regions are again maximally correlated with the SAL network, while the ventral putamen seed regions are again maximally correlated with the CG-OP network. The VS seed regions also show strong, positive correlation with the CG-OP network showing the pattern is preferential not selective. Data plotted here are a re-visualization of (i.e., not independent from) the data plotted in Figures 9-15. Bars show mean correlation (*z*(*r*)) in each network, error bars indicate standard error of the mean adjusted for within-subject variance (Cousineau 2005).

Across all 15 participants, the correlation pattern produced by the ventral putamen seed regions were again substantially different than the pattern produced by the VS seed regions. First, the ventral putamen seed regions were most strongly correlated with the CG-OP network in DLPFC (Figure 16, top right) and across the full extent of the cerebral cortex (Figure 16, bottom right). Second, correlations between the ventral putamen seed regions and FPN-A were indistinguishable from zero.

Taken together, these results demonstrate that distinct patterns of cortical connectivity for VS and ventral putamen seed regions are found consistently across participants in relation to the SAL and CG-OP networks, and further that neither the VS nor ventral putamen are positively correlated with FPN-A, a network implicated in cognitive control.

### Ventral Striatum and Ventral Putamen Seed Regions Are Anatomically Adjacent

Idiosyncratic anatomical variability was present across individuals in the locations of the striatal seed regions associated with the SAL and CG-OP networks. To visualize the relative placements of seed regions within the striatum, the positions of the seed regions from all 17 participants were plotted together. The seed regions that recapitulated the cortical SAL network were predominantly localized to the rostral and medial portions of the ventral striatum, proximal to the caudate (Figure 17, pink circles). Conversely, the seed regions that recapitulated the CG-OP network were localized to ventral portions of the putamen, caudal and lateral to the VS seed regions (Figure 17, purple circles).

**Figure 17.**
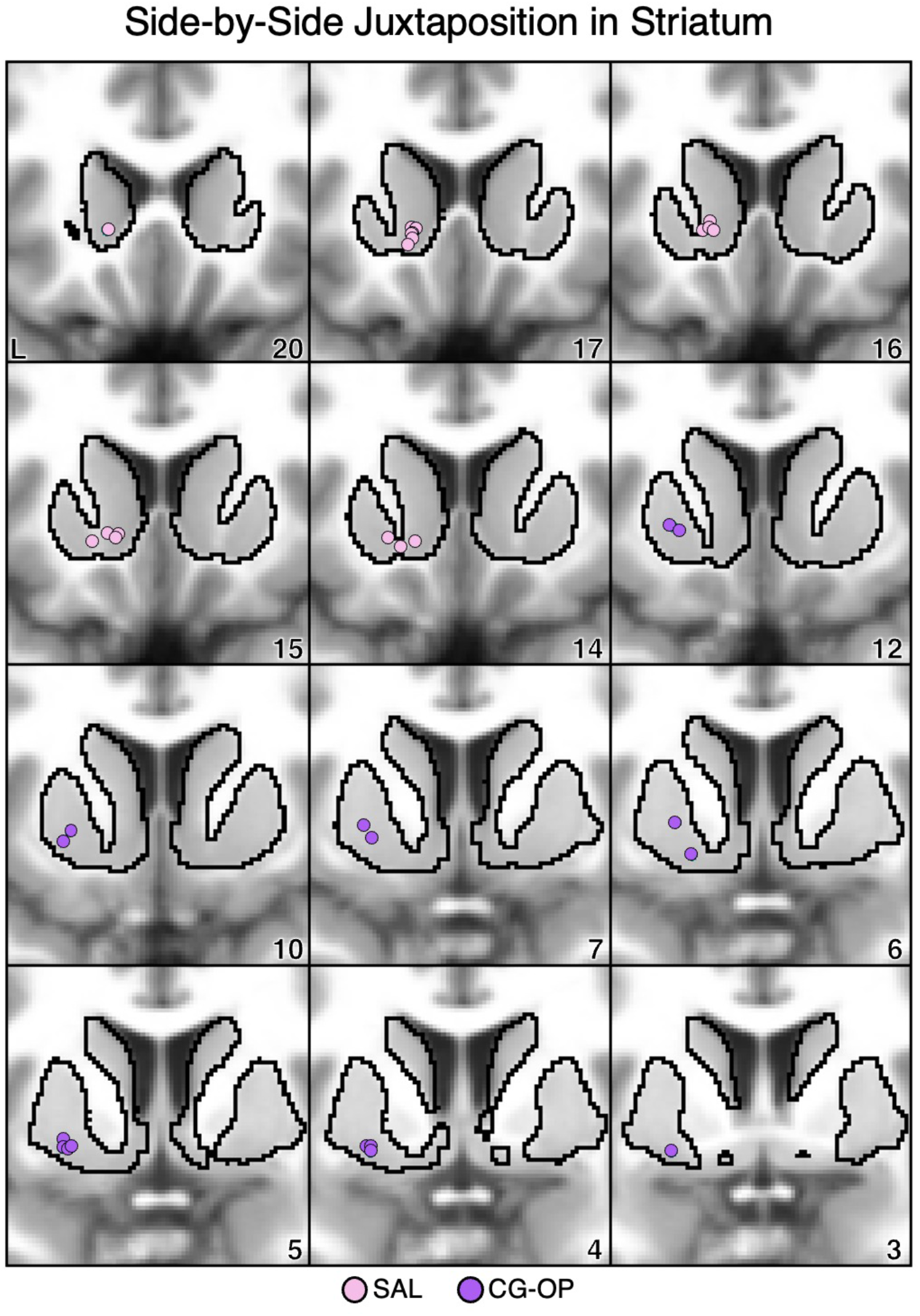
Ventral Striatum (VS) and Ventral Putamen Seed Regions are Anatomically Juxtaposed. Coronal views of the striatum show seed regions that are correlated with the SAL network (pink circles) are rostral and medial to seed regions that are correlated with the CG-OP network (purple circles). Each dot represents an individual participant (N=17, including S1 and S2, and P1-P15). The T1-weighted anatomical image backdrop is the average from all 17 participants transformed to the MNI atlas space. The black outlines indicate the average boundary of the striatum in MNI atlas space. L = left. Coordinates in the bottom right of the left panels indicate the y coordinates (in mm).

## Discussion

The striatum receives projections from the cerebral cortex with separation between motor, association, and limbic zones. Here we demonstrate using fcMRI that the VS, which includes the NAc, is preferentially correlated with a distributed cortical network known in the human literature as the SAL network, including a specific region in DLPFC. Moreover, the region of the VS associated with the SAL network is distinct from an anatomically adjacent region in the ventral portion of the putamen that is preferentially correlated with the CG-OP network, including a distinct region in DLPFC. Neither of these ventral striatal regions correlates with regions implicated in cognitive control including in the DLPFC. We discuss the implications of these findings for understanding striatal organization and for targeted neuromodulation.

### The Ventral Striatum is Preferentially Correlated with the Salience Network

Broad separation between motor, association, and limbic zones was established based on anatomical tracing studies in monkeys (Alexander, DeLong, and Strick 1986; Haber and McFarland 1999; Heimer 1978) and confirmed in humans using neuroimaging (Choi, Yeo, and Buckner 2012; Di Martino et al. 2008; Greene et al. 2014; Jarbo and Verstynen 2015; Marquand, Haak, and Beckmann 2017; Morris et al. 2016). Recent precision estimates in the human have revealed that there is considerable structure beyond these broad distinctions even within domains. For example, the association zone of the striatum, specifically the caudate extending into the dorsal putamen, maintains segregation (or partial segregation) of distinct cortical networks that are involved in cognition (Gordon et al. 2022; Kosakowski et al. 2024). In the present work we employed within-individual precision neuroimaging to show that the VS participates in the SAL network which can be distinguished from a juxtaposed region in the ventral putamen linked to the CG-OP network.

The SAL and CG-OP networks are distributed brain-wide networks. While their contributions to information processing are complex and debated, the two networks are linked to perceiving and reacting to changing or unexpected events (Dosenbach et al. 2006; Seeley et al. 2007; for recent discussion see Gordon et al. 2023; Seeley 2019). Both networks are adjacent to one another across multiple regions of cortex leading to confusion especially in group-averaged studies (Seeley 2019). Within-individual precision fcMRI estimates employed here (and elsewhere) can separate the two networks. Precision estimates of the SAL and CG-OP networks have previously suggested that the two networks are linked to the striatum at or around the VS extending into the putamen (Gordon et al. 2023; Lynch et al. 2024). Our results build on these prior findings and indicate that the SAL network is specifically linked to a region in the VS at and around the NAc core, and further that the CG-OP network is linked to an adjacent region in the ventral putamen. These finding suggest the two separate, but adjacent, cortical networks maintain their anatomical separation within and near to the VS.

Lynch et al. (2024) demonstrated that the SAL network includes the VS (see their Figure 1e). Our findings are fully consistent with this observation. In S1 and S2, the patterns of cortical correlations with seed regions in the VS recapitulated the full distributed extent of the SAL network, including canonical regions in the anterior insula and cingulate, as well as a specific region of DLPFC. The patterns were stable across independent datasets from the same individuals (Figures 1 and 2) and generalized across 15 additional individuals (Figures 9-11). Quantitative analyses in independent data in S1 and S2 further revealed the VS seed region is preferentially correlated with the SAL network more than any other network in association cortex (Figure 6). Thus, by all analyses the VS is associated with the cortical network known as the SAL network in the human literature. Several additional details are notable.

First, the region of the VS associated with the SAL network could be anatomically double dissociated from a juxtaposed region that fell within the ventral putamen. The region in the ventral putamen was positioned near to, but distinct from, the topographic representation of the motor effector zones that have been previously mapped in the putamen of S1 and S2 (Kosakowski et al. 2024). This is a particularly interesting arrangement because the cortical network linked to the ventral putamen is the CG-OP network, a network that surrounds the motor and somatosensory areas in the cortex along pre- and post-central gyri, including components that create discontinuities in the motor map (so called ‘inter-effector’ regions; Gordon et al. 2023). That is, striatal regions associated with the CG-OP network fall in-between the VS, which is linked to the SAL network, and the region of the putamen that is associated with topographically mapped motor representations. These findings suggest that the organization of the striatum is anatomically specific and recapitulates broad organizational adjacencies present in the cerebral cortex.

Second, the predominant cortical network linked to the VS is the SAL network and not the ‘Default’ network as we and others previously hypothesized (e.g., Choi et al. 2012; Di Martino et al. 2008). As raised in the introduction, in group-averaged data, seed regions placed along the MPFC can yield a local correlation pattern that blurs across the adjacent (narrow) white-matter tract into the VS. A control analysis in the present within-individual data revealed that seed regions placed within the MPFC, specifically in the location at or near sgACC, did not yield correlations that extended into the VS (Figure 7). Rather, a correlation pattern in the cerebral cortex emerged that robustly included the distributed regions linked to the cortical network known as DN-A (Braga and Buckner 2017; Braga et al. 2019; Du et al. 2024; see also Angeli et al. 2023; Reznik et al. 2023; Zhang et al. 2021). These observations again suggest anatomical specifically in the VS and further clarify how the present results and those of Lynch et al. (2024) can be reconciled with prior estimates of the organization of the VS. We suspect earlier work proposing a prominent association between the VS and the ‘Default’ network may have been an artifact of spatial blurring. The present precision estimates converge with Lynch et al. (2024) to show that the VS is preferentially correlated with the SAL network.

### The Ventral Striatum is Linked to the Dorsolateral Prefrontal Cortex

An intriguing finding is that specific regions of the DLPFC are components of the distributed cortical networks linked to the VS and ventral putamen. This robust finding is observed in every participant examined and has both conceptual implications for the study of PFC function as well as translational implications for therapeutic strategies.

The PFC is well established to play a role in executive function including supporting cognitive control (Badre and D’Esposito 2009; Duncan, 2010; Duncan and Owen 2000; Friedman and Robbins 2022; Miller 2000; Miller and Cohen 2001). Recent analyses using within-individual precision methods have revealed that there are regions within PFC that are functionally dissociable from those participating in domain-flexible aspects of cognitive control (e.g., DiNicola, Sun, and Buckner 2023; Du et al. 2024; Fedorenko et al. 2012; see also Fedorenko and Thompson-Schill 2014; Somers et al. 2021). Fedorenko and colleagues (2012), in a series of foundational studies, provided strong evidence that ventrolateral PFC regions specialized for language are near to, but dissociable from, adjacent regions that respond broadly to increases in cognitive effort (see also Braga et al., 2020; Du et al., 2024; Fedorenko and Thompson-Schill 2014). DiNicola et al. (2023) further demonstrated that even within DLPFC there are dissociable regions that participate in processing spatial information that are distinct from nearby regions involved in domain-flexible aspects of working memory. The specific regions of PFC that are preferential for information processing domains appear to gain their properties through the distributed networks that they are situated within. Here we demonstrate that there are distinct regions in DLPFC embedded within the SAL and CG-OP networks that are functionally connected to the VS and ventral putamen, respectively. This observation adds to our understanding of the heterogeneity of PFC and provides potential targets for neuromodulation.

The pattern of correlations with the VS and ventral putamen, in every participant, yielded a correlated region (or sometimes multiple regions) in DLPFC that was distinct from regions implicated in cognitive control (Figures 5, 15). The specific regions of DLPFC linked to the VS are at or near the site targeted by Transcranial Magnetic Stimulation (TMS) treatment of MDD (for review of TMS treatment of MDD see Perera et al. 2016). Standard treatment protocols target left DLPFC 4-5 cm anterior to the location of motor cortex (Pascual-Leone et al. 1996), the left F3 location in the 10-20 coordinate system (Herwig, Satrapi, and Scho nfeldt-Lecuona 2003; Okamoto et al. 2004), or the Beam F3 location (Beam et al. 2009). In an early group-based examination of TMS target efficacy, Fox et al. (2012) proposed that the most effective site for TMS therapy in MDD is the specific DLPFC region anti-correlated (negatively correlated) with the sgACC. In recent clinical trials that have paired dense, repeated sessions of intermittent theta burst stimulation (iTBS) with within-individual estimation of the anti-correlated sgACC target, clinical efficacy has improved (Cole et al. 2020; 2022).

While the mechanisms of the treatment’s improvement are unknown (e.g., increased dosing, shift in general location, individualized targeting), it is intriguing that the most effective DLPFC targets fall quite close to the DLPFC region coupled to the VS. In a recent quantitative analysis of the brain networks targeted using the left F3 location or those targeted using the sgACC anti-correlation approach, the achieved DLPFC target in both cases involved the SAL network in most participants (Sun et al. 2024). That is, while the VS-correlated regions of the DLPFC make up only a small portion of the PFC, they are preferentially targeted in TMS treatment for MDD raising the possibility that the mechanism-of-action is related to stimulation of the VS or circuitry linked to the VS.

### Open Questions and Limitations

Estimates of connectivity in humans using fcMRI are based on an indirect proxy of anatomical connectivity (Buckner, Krienen, and Yeo 2013; Fox and Raichle 2007; Murphy et al. 2013; Power, Schlaggar, and Petersen 2014; Smith et al. 2013; Van Dijk et al. 2010). As such, contrasting network estimates from studies in humans with direct anatomical connectivity in monkeys is informative to interpret results. Such comparisons reveal good correspondence between anatomical projection patterns in the putamen in relation to motor topography as well as broad correspondence for network patterns associated with the caudate (Choi et al. 2012). However, while the cortical VS correlation pattern is reproducible across participants and laboratories, examining the details of the anatomical connectivity of the VS in the monkey leaves unresolved how the present indirect estimates of network connectivity in the human relate to anatomical connectivity in the monkey.

Specifically, anterograde injections in DLPFC do not yield prominent label in the VS (e.g., Case 1 in Selemon and Goldman-Rakic 1985; Case 5 in Yeterian and Pandya 1991; Case OM38 in Ferry et al. 2000; Case 131 in Calzavara, Mailly, and Haber 2007) nor do retrograde injections to the VS yield a pattern that consistently includes DLPFC (e.g., Case MR28WGA in Choi, Ding, and Haber 2017; see also VSv in their Table 2). DLPFC tracer injections yield robust projections to the caudate extending across the internal capsule into the putamen (for review see Haber 2016). Injections to the dorsal extent of the VS (VSd) reveal dense projections from widely distributed cortical association regions including the DLPFC (e.g., Cases MN38LY and MN40LY in Choi, Ding, and Haber 2017). Thus, one possibility is that the observed connectivity in the human reflects VS connectivity in the NAc core extending into a homologue of VSd, and patterns linked to the NAc shell are not fully captured in our analyses. Another possibility is that the correlated network linked to the VS measured with fcMRI reflects polysynaptic projections. Higher resolution data in humans and more comprehensive mapping of association zones in monkeys may provide further insights.

Another limitation is that the present study explored network connectivity and did not directly address functional response properties. Having established that the VS and ventral putamen are linked to the SAL and CG-OP networks, it will be important for future research to investigate functions related to reward and motivation in relation to these findings, including the specific SAL network regions in the DLPFC that are linked to the VS and are at or near current neuromodulation targets for MDD. SAL network function has most often been explored in relation to transient responses to changing and unexpected events, while VS function has most often been explored in relation to the anticipation and receipt of reward. The discovery that the VS is linked to the SAL network raises the question of how these two ways of probing the brain’s response to dynamic, changing events are related.

## Conclusions

The VS is correlated in an anatomically specific manner with the distributed cortical network known in the human literature as the SAL network. The region of the VS linked to the SAL network can be distinguished from an adjacent region in the ventral putamen that is associated with the CG-OP network. The cortical networks linked to both the VS and ventral putamen include specific regions in the DLPFC which may be viable targets for neuromodulation.

## Acknowledgments

We thank the Harvard Center for Brain Science neuroimaging core and FAS Division of Research Computing. We thank T. O’Keefe for assisting in optimization of data processing, R. Mair for MRI physics support, and P. Angeli, L. DiNicola, W. Sun, A. Billot, V. Tripathi, and A. Xue for discussion and technical assistance. The multi-band EPI sequence was provided by the Center for Magnetic Resonance Research (CMRR) at the University of Minnesota.

## Footnotes

1 The label FPN-A is used for this network (as used in Xue et al. 2021 and Du et al. 2024). In some prior reports, this network was referred to as FPN-B (e.g., Braga et al. 2020).

